# tRF-3021a Supports Invasiveness, Apoptosis Resistance, and Global Protein Synthesis in Cancer Cell Lines

**DOI:** 10.64898/2026.01.26.701835

**Authors:** Esmaeili Fatemeh, Kumarjeet Banerjee, Ajay Chatrath, Divya Sahu, Yoshiyuki Shibata, Shekhar Saha, Pankaj Kumar, Zhangli Su, Anindya Dutta

**Affiliations:** Department of Genetics, University of Alabama at Birmingham; Department of Biochemistry and Molecular Genetics, University of Virginia

**Author notes:** co-corresponding authors: Dr. Anindya Dutta, Dr. Zhangli Su, Department of Genetics, University of Alabama at Birmingham. Conflict of Interest: The authors declare no conflicts of interest.

## Abstract

tRNA-derived fragments (tRFs) are a relatively recently discovered class of small RNAs implicated in gene-regulatory processes in diverse biological contexts but there have been very few reports of a clear phenotypic role of these small RNAs in cancer progression. To select tRFs that should receive priority for mechanistic experiments, we analyzed small RNA-seq data from The Cancer Genome Atlas (TCGA) and found that high expression of three 3′ tRFs (tRF-3a), tRF-3009a, tRF-3021a or tRF-3030a, is significantly associated with poor overall survival in low-grade glioma (LGG). In glioblastoma cells, tRF-3009a, tRF-3021a and tRF-3030a enhance cell invasion and migration but tRF-3021a was uniquely required for cell proliferation and suppression of apoptosis. Interestingly, tRF-3021a knockdown decreases global protein synthesis prior to and independent of apoptosis in a number of cancer cell lines, both from and outside the glioblastoma lineage. RNA-seq reveals that the results cannot be explained by microRNA-like functions of tRF-3021a. These data indicate that tRF-3021a supports cancer cell survival and particularly protein synthesis while promoting cellular invasion and migration and suggests that strategies to downregulate this short RNA could be a potential therapeutic approach in cancers.

**Implication:** tRF-3021a promotes malignant cell phenotypes, sustains global protein synthesis and prevents spontaneous apoptosis, motivating efforts to evaluate it as a biomarker and therapeutic target.

## Introduction

In addition to well-studied small noncoding RNA (sncRNA) regulators, such as microRNAs (miRNAs), small interfering RNAs (siRNAs), and PIWI-interacting RNAs (piRNAs), which regulate gene expression through association with Argonaute family proteins (AGO-clade Argonautes for miRNAs/siRNAs and PIWI-clade Argonautes for piRNAs) to form RNA-induced silencing complexes (RISC) [1, 2], studies over the last fifteen years have identified additional sncRNAs derived from either mature or precursor tRNAs [3–6]. Collectively termed tRNA-derived small RNAs (tsRNAs), these molecules are commonly grouped into (a) tRNA halves (∼31–40 nt), including 5′ halves (from the 5′ end to the anticodon loop) and 3′ halves (from the anticodon loop to the 3′ CCA end), which are often produced from mature tRNAs under stress conditions, and (b) shorter tRNA-derived fragments (tRFs). tRFs include tRF-1 (derived from the 3′ trailer sequence of precursor tRNAs), tRF-5 (derived from the 5′ end of mature tRNAs and often subdivided into tRF-5a [∼14–16 nt], tRF-5b [∼22–24 nt], and tRF-5c [∼28–30 nt]), and tRF-3 (derived from the 3′ end of mature tRNAs, spanning the TΨC loop to the 3′ CCA end, and often subdivided into tRF-3a [∼18 nt] and tRF-3b [∼22 nt]). In addition, i-tRFs and tRF-2 species arise from internal tRNA regions and the anticodon loop, respectively [7–9].

The precise nucleases responsible for generating each class of tRFs remain largely unknown. For tRNA halves, angiogenin has been shown to be required at least for 5′ tRNA-His(GTG) and 3′ tRNA-Asp(GTC), but angiogenin knockout does not abolish the production of other tRNA halves [10]. In contrast, tRF-1 generation depends on RNase Z (ELAC2) [5]. For tRF-5 and tRF-3 species, some studies report DICER dependence, whereas others show these fragments can be produced independently of DICER [3, 4, 11–13]. Their conservation across all three domains of life, along with reproducible cleavage sites, defined sizes, and high abundance, argues against nonspecific degradation.

tRFs and tRNA halves have both physiological and pathophysiological roles and can act through multiple modes of action, including direct binding to RNAs or proteins such as RNA-binding proteins (RBPs) [9]. They have been shown to regulate cell proliferation and cell cycle progression [5, 14], and some function in a microRNA-like manner by recognizing target mRNAs through seed-region rules and repressing translation [12, 13, 15, 16]. In addition, these RNAs can bind RBPs and modulate their activity [17–19], promote nuclear localization of proteins through direct interaction [20], and participate in epigenetic and chromatin regulation [9, 21]. tRFs and tRNA halves have also been implicated in viral infection [22–24], and in translational control, with reports of both global translation inhibition [25] and enhanced translation via effects on ribosome biogenesis [26]. Consistent with these diverse functions, these RNAs are implicated in multiple diseases, including cancer, where they can act as tumor suppressors or oncogenes depending on context [9, 18]. Since microRNAs and other noncoding RNAs have been widely studied in cancers, we sought evidence that clearly implicates tRFs in cancer cell progression. A screen for cancers where levels of any of the tRFs predict outcome revealed that overexpression of three specific tRFs predicts poor outcome in lower grade gliomas (LGG). Following up on this, we discovered that these tRFs promote cancer cell invasion and migration in multiple cancer cell lineages, but most interesting, one of them, tRF-3021a is unique in being required for normal protein synthesis, such that its inhibition rapidly leads to cancer cell apoptosis.

## Materials and Methods

### Cell Culture and Treatment

The human GBM cell line U-87 MG (ATCC HTB-14, RRID:CVCL_0022) was purchased from American Type Culture Collection (ATCC). U-251 MG (RRID:CVCL_0021) and normal human astrocytes (RRID:CVCL_E3G4) were generous gifts from Roger Abounader lab (University of Virginia). HCT116 (RRID:CVCL_0291) was a gift from Bert Vogelstein. MCF7 (HTB-22 ™, RRID:CVCL_0031), and U-2 OS (HTB-96 ™, RRID:CVCL_0042) cells were purchased from ATCC. Cells were maintained at 37 °C in a humidified incubator with 5% CO₂ and cultured in DMEM (Thermo Fisher Scientific, 31-053-028) supplemented with 10% fetal bovine serum (FBS, Thermo Fisher Scientific, 10437028) and 1% penicillin/streptomycin, except HCT116, which was maintained in McCoy’s 5A medium.

For chemical treatment, cells were treated with the pan-caspase inhibitor z-VAD-fmk (InvivoGen, tlrl-vad) at a final concentration of 20 µM for 48 h. To maintain inhibition, fresh z-VAD-fmk was added again at 24 h.

### Transfections

For tRF knockdown (KD), cells were reverse-transfected with mixmer/gapmer LNA antisense oligonucleotides or a sequence-scrambled negative control LNA (Integrated DNA Technologies, IDT) using Lipofectamine RNAiMAX (Thermo Fisher Scientific, 13778075) according to the manufacturer’s instructions. LNA sequences were as follows: Control LNA: +GTA+CG+CG+GAA+TA+C+TT+C; tRF-3009a LNA: TGGTACCA+GG+AG+TGG+GG+T; tRF-3021a LNA: T+GG+TGG+AG+ATGC+CGGG+GA; tRF-3030a LNA: T+GG+TCC+TT+CG+AG+CCG+GA+A; tRF-3021a gapmer LNA: +G+G+T+GGAGATGCCG+G+G+G+A; α 5’ LNA: G+CT+CTA+CC+ACTG+AGCT+AC; α Anticodon LNA: C+GG+GAC+CT+CATA+CATG+CA; PM1 LNA: T+GG+TGG+AG+GTGC+CGGG+GA; PM3 LNA: T+GG+TGG+AG+GT+AC+TGGG+GA; hsa-miR-7-5p LNA: A+AC+AAC+AA+AATC+ACTA+GTC+TTC+CA; hsa-miR-9-5p LNA: T+CA+TAC+AG+CTAG+ATAA+CCA+AA+GA. Unless otherwise indicated, LNAs were used at 30 nM in antibiotic-free DMEM supplemented with 10% FBS, and cells were harvested 2–48 h post-transfection for downstream assays.

For overexpression, cells were reverse-transfected with synthetic RNA mimics or a scrambled control mimic (IDT) using Lipofectamine RNAiMAX according to the manufacturer’s instructions. Mimic sequences (all containing 2′-O-methyl–modified ribose and 5′ phosphorylation) were: ssGL2_2ome: /5Phos/mCrGmUrCmGrCmGrGmArAmUrAmCrUmUrCmGrAmUrU; 3009a_2ome: /5Phos/mArCmCrCmCrAmCrUmCrCmUrGmGrUmArCmCrA; 3021a_2ome: /5Phos/mUrCmCrCmCrGmGrCmArUmCrUmCrCmArCmCrA; 3030a_2ome: /5Phos/mUrUmCrCmGrGmCrUmCrGmArAmGrGmArCmCrA; tRF-3021a mimic Mutant 1: /5Phos/mGrAmArCmCrGmGrCmArUmCrUmCrCmArCmCrA; tRF-3021a mimic Mutant 2: /5Phos/mUrCmCrUmUrAmGrCmArUmCrUmCrCmArCmCrA; tRF-3021a mimic Mutant 3: /5Phos/mUrCmCrCmCrGmUrAmGrCmCrUmCrCmArCmCrA; tRF-3021a mimic Mutant 4: /5Phos/mUrCmCrCmCrGmGrCmArUmArGmArAmArCmCrA; tRF-3021a mimic Mutant 5: /5Phos/mUrCmCrCmCrGmGrCmArUmCrUmCrCmGrAmArG. Unless otherwise specified, mimics were used at 50 nM.

### Cell proliferation assay

Cells were seeded in 96-well plates (10³ cells/well). At 24, 48, and 72 h, MTT (5 mg/mL stock, Promega, G4100) was added to a final 0.5 mg/mL and incubated 4 h at 37 °C. Formazan crystals were solubilized by adding 100 µL stop/solubilization solution per well, and absorbance was measured at 570 nm using a BioTek Synergy H1 microplate reader (Agilent/BioTek, RRID:SCR_019748). Data were collected using Gen5 software (RRID:SCR_017317). Values were normalized to the mean absorbance of control wells.

### Cell migration and invasion assays

Cell migration and invasion were assessed using Transwell inserts in 24-well format. Invasion assays were performed using Corning BioCoat Matrigel Invasion Chambers (08-774-122), whereas migration assays were performed using Falcon Permeable Supports with 8.0 µm PET membranes (Sigma, CLS353097). For migration, inserts were used uncoated, while for invasion, Matrigel-coated inserts were used. For the experiments shown in Fig. 2A–F, cells were first reverse-transfected in 6-well plates. Twenty-four hours after transfection, cells were reseeded into Transwell inserts and allowed to migrate or invade for 18–24 h. Thus, cells were fixed and analyzed 42–48 h after the initial transfection. For the 12 h Transwell assays shown in Supplementary Fig. 1I-K, cells were reverse-transfected directly in the Transwell inserts and fixed 12 h after transfection. DMEM + 20% FBS (800 µL) was placed in the lower chamber as a chemoattractant. At the indicated endpoint, non-migrated or non-invaded cells were removed from the upper side of the membrane. Membranes were fixed in 4% paraformaldehyde, stained with 0.5% crystal violet, and imaged. Cells on the lower surface were manually counted in ≥3 random fields per insert at 10× magnification.

**Figure 1.**
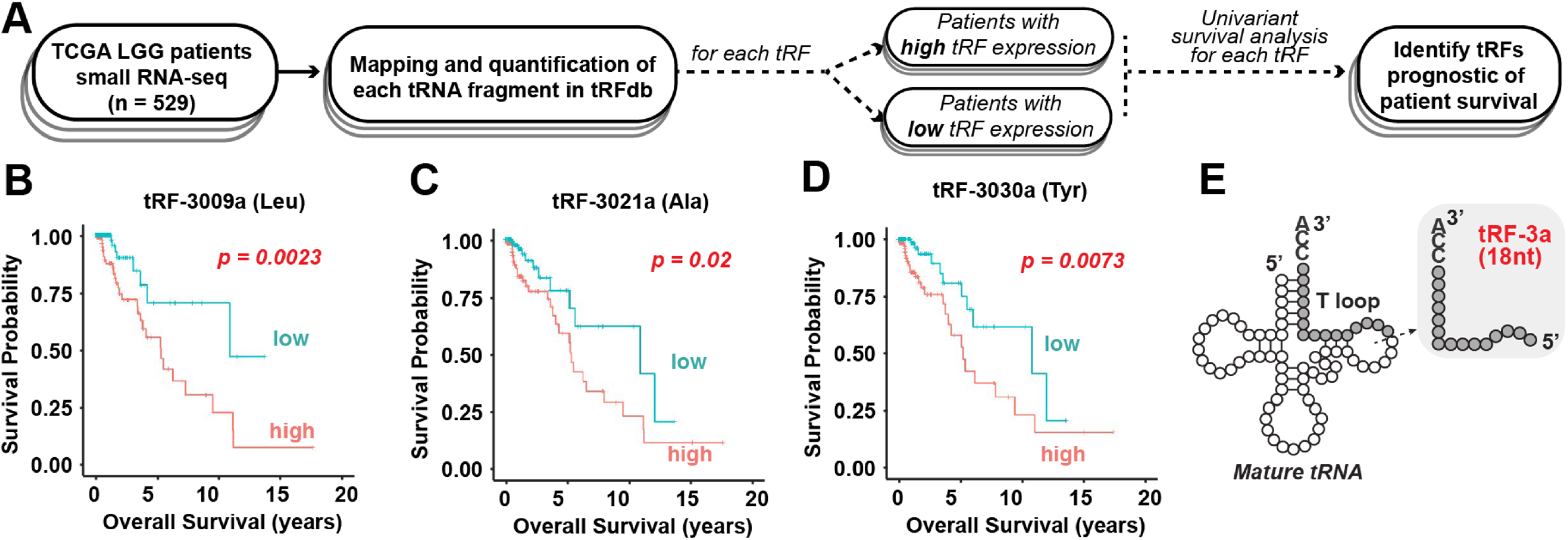
High expression of tRF-3s is associated with poor prognosis in LGG patients. (A) Workflow for identifying prognostic tRNA-derived fragments (tRFs) in TCGA lower-grade glioma (LGG) patients using small RNA-seq data and overall survival. (B–D) Kaplan–Meier survival curves for LGG patients stratified by expression of tRF-3009a (B), tRF-3021a (C) and tRF-3030a (D). Patients were divided into the top third (high expression, red line) and bottom third (low expression, blue line) of tRF expression. (tRF-3009a, p = 0.0023, tRF-3021a, p = 0.02, tRF-3030a, p = 0.0073). (E) Secondary structure of the parental mature tRNAs showing the position of the 18-nt tRF-3a derived from the 3′ end of mature tRNA (TψC loop to CCA).

**Figure 2.**
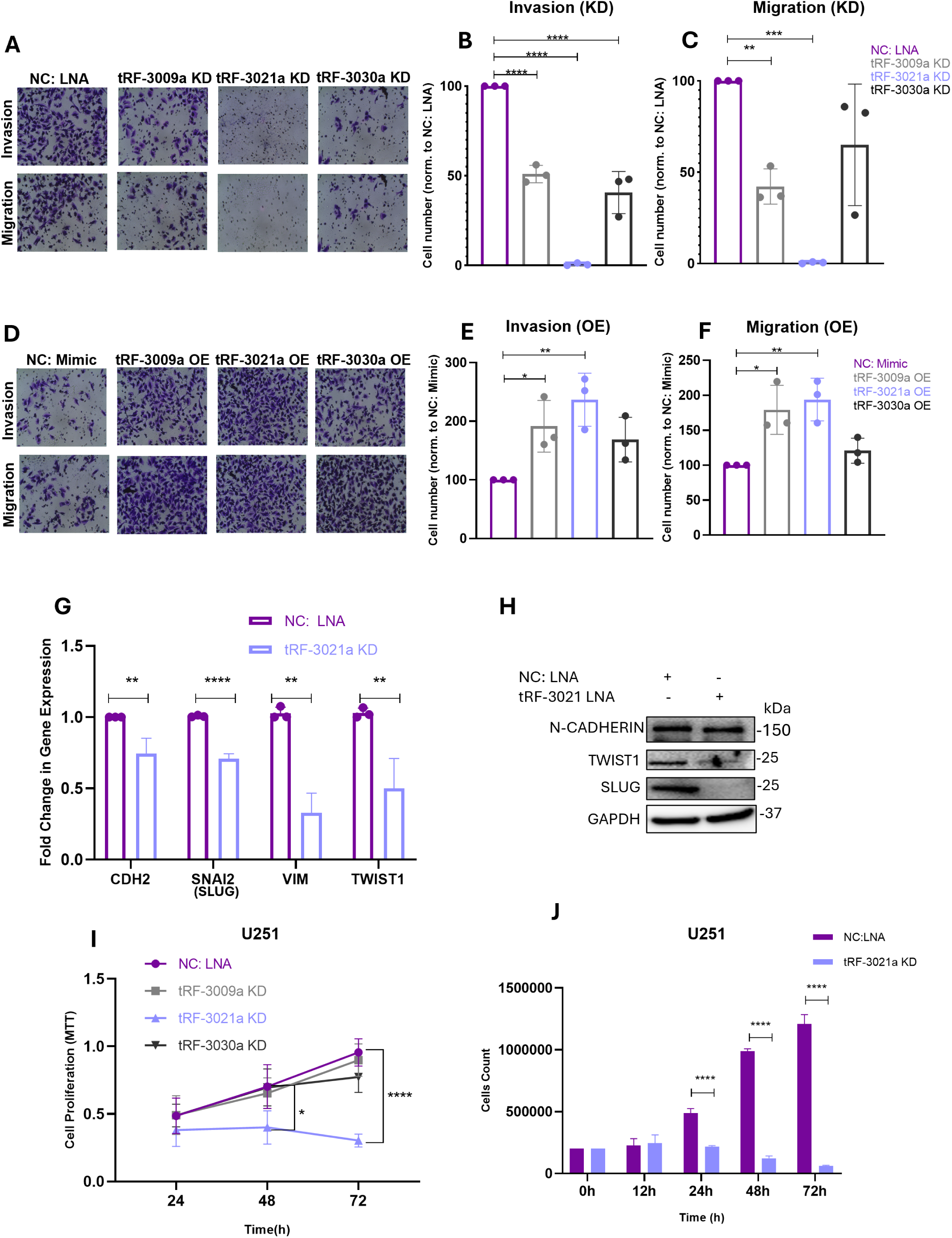
tRF-3 fragments regulate invasion, migration, and proliferation of GBM cells. (A) Representative Transwell images of invasion and migration of U251 cells after LNA-mediated knockdown (KD) of tRF-3009a, tRF-3021a, or tRF-3030a compared with negative-control (NC) LNA. (B–C) Quantification of invasion (B) and migration (C) after tRF-3 KD. Cell numbers were normalized to NC LNA, which was set to 100%. (D) Representative Transwell images of invasion and migration of U251 cells after mimic-mediated overexpression (OE) of tRF-3009a, tRF-3021a, or tRF-3030a compared with NC mimic. (E–F) Quantification of invasion (E) and migration (F) after tRF-3 OE. Cell numbers were normalized to NC mimic, which was set to 100%. (G) qRT-PCR analysis of invasive-state transcripts, including *CDH2*, *SNAI2* (Slug), *VIM*, and *TWIST1*, in U251 cells after tRF-3021a KD compared with NC LNA. Expression was normalized to *ACTB* and shown relative to NC LNA. (H) Immunoblot analysis of N-cadherin, TWIST1, and SLUG in U251 cells after tRF-3021a KD compared with NC LNA. GAPDH was used as a loading control. (I) U251 cell proliferation measured by MTT assay at 24, 48, and 72 h after KD of tRF-3009a, tRF-3021a, or tRF-3030a compared with NC LNA. (J) Direct cell counting of U251 cells after NC LNA or tRF-3021a LNA transfection at 0, 12, 24, 48, and 72 h. Data are presented as mean ± SD from independent biological replicates unless otherwise indicated. Statistical significance was determined by one-way ANOVA with Dunnett’s multiple-comparisons test for invasion and migration KD and OE experiments, unpaired two-tailed t-test for qRT-PCR comparisons, and two-way ANOVA with multiple-comparisons testing for MTT and cell-counting time-course experiments. ns, not significant, *P < 0.05, **P < 0.01, ***P < 0.001, ****P < 0.0001.

### Annexin V/PI staining

Cells were trypsinized, washed with PBS, and processed according to the manufacturer’s instructions using the Dead Cell Apoptosis Kit with Annexin V for Flow Cytometry (Thermo Fisher/Life Technologies, V13242). Samples were analyzed on a BD LSRFortessa flow cytometer (RRID:SCR_018655) at the UAB Flow Comprehensive Flow Cytometry Core. The percentages of Annexin V⁺/PI⁻ (early apoptotic) and Annexin V⁺/PI⁺ (late apoptotic) cells were quantified using FlowJo (RRID:SCR_008520).

### Western blotting and puromycin pulse labeling

Cells were lysed in RIPA buffer (50 mM Tris-HCl, pH 8.0, 150 mM NaCl, 1.0% NP-40, 0.5% sodium deoxycholate and 0.1% SDS). Equal amounts of protein were resolved by 10% SDS-PAGE, transferred to PVDF membranes using Bio-Rad Trans-Blot semi-dry transfer machine and immunoblotted with primary antibodies (all 1:1000) against GAPDH (Santa Cruz Biotechnology, sc-32233, RRID:AB_627679), TWIST (Santa Cruz Biotechnology, sc-15393, RRID:AB_2211738), N-cadherin (Cell Signaling Technology, 13116T, RRID:AB_2687616), Slug (Cell Signaling Technology, 9585, RRID:AB_2239535), PARP (Cell Signaling Technology, 9542, RRID:AB_2160739), cleaved PARP (Asp214) (Cell Signaling Technology, 5625T, RRID:AB_10699459), phospho-Histone H2A.X (Ser139) (Cell Signaling Technology, 2577, RRID:AB_2118010), total eIF2α (Cell Signaling Technology, 9722S, RRID:AB_2230924), phospho-eIF2α (Ser51) (Cell Signaling Technology, 9721S, RRID:AB_330951), total 4E-BP1 (Cell Signaling Technology, 9452, RRID:AB_331692), phospho-4E-BP1 (Thr37/46) (Cell Signaling Technology, 2855T, RRID:AB_560835), total p70 S6 kinase (Cell Signaling Technology, 9202, RRID:AB_331676), and phospho-p70 S6 kinase (Thr389) (Cell Signaling Technology, 9234, RRID:AB_2269803). GAPDH was used as the loading control. Proteins were detected using an ECL Immobilon Western Chemiluminescent HRP Substrate detection kit (Millipore, WBKLS0500) and imaged with ChemiDoc MP imaging system (BIO-RAD, RRID:SCR_019037). For puromycin labeling, culture medium was replaced with fresh complete medium containing puromycin (10 µg/mL), and cells were incubated for 15 min at 37 °C before lysis as described above. Puromycin incorporation was detected using anti-puromycin antibody (Kerafast, Kf-Ab02366-1.1 (EQ0001), mouse IgG1, 1:1000, RRID:AB_2620162). Anti-rabbit IgG, HRP-linked Antibody (Cell Signaling Technology, 7074S, RRID:AB_2099233) and Anti-mouse IgG, HRP-linked Antibody (Cell Signaling Technology, 7076S, RRID:AB_330924) were used with 1:5000 dilution as secondary antibodies.

### L-homopropargylglycine (HPG) assay

Briefly, U251 cells were transfected with 30 nM tRF-3021a LNA antisense oligonucleotide or sequence-scrambled negative control. Twenty-four hours after transfection, cells were incubated in methionine-free growth medium (Thermo Fisher Scientific, 21013024) for 60 min at 37°C. Cells were then incubated with 100 µM L-homopropargylglycine (HPG, Vector Laboratories, CCT-1067-25) at 37°C for 60 min, followed by PBS washing and fixation with 4% paraformaldehyde at room temperature for 15 min. Cells were washed with PBS and permeabilized with PBS containing 0.3% Triton X-100 for 10 min at room temperature. HPG incorporation was detected using a freshly prepared click chemistry reaction cocktail containing PBS, 1 mM CuSO₄, 10 mM THPTA ligand (Vector Laboratories, CCT-1010-100), 5 µM AZDye 594 azide (Vector Laboratories, CCT-1295-1), and sodium ascorbate. Sodium ascorbate was added last, and the reaction cocktail was added immediately to the cells. Cells were incubated with the click reaction cocktail for 30 min at room temperature protected from light, washed three times with PBS, and counterstained with DAPI for 1 min at room temperature. Fluorescence images were acquired using a Zeiss laser-scanning confocal microscope (RRID:SCR_017377).

### Dual luciferase assay

To generate the tRF-3021a complementary reporter construct, sense and antisense oligonucleotides containing the sequence perfectly complementary to tRF-3021a were ordered from IDT with 5′ overhangs compatible with XhoI- and NotI-digested psiCHECK-2 vector. The oligonucleotides were phosphorylated and annealed using T4 DNA Ligase Buffer (NEB, B0202S) and T4 Polynucleotide Kinase (T4 PNK, NEB, M0201S), according to the manufacturer’s protocol. The empty psiCHECK-2 vector was digested with XhoI (NEB, R0146S) and NotI (NEB, R0189S), and the annealed double-stranded oligonucleotide insert was ligated into the digested vector using 10× T4 DNA Ligase Buffer and T4 DNA Ligase (NEB, M0202S), according to the manufacturer’s protocol.

tRF-target luciferase assays to assess tRF activity were performed in 24-well plates. 50 nM of tRF mimic or a scrambled control mimic or 30 nM of LNA antisense oligonucleotides and 500 ng of psi-CHECK2 reporter (Promega, C8021) vector with perfect complementary sites to tRF-3021a in the 3′UTR of renilla luciferase gene were co-transfected into U251 cells using Lipofectamine 2000 (Thermo Fisher Scientific, 11668027) as a transfection reagent. The cells were lysed in 100 µL 1× passive lysis buffer from the Dual-Luciferase Reporter Assay System (Promega, E1980) after 48 h of transfection. The luciferase signals were measured using 20 µL of the lysate following the manufacturer’s protocol. The renilla luciferase levels were normalized to firefly luciferase levels. Luminescence was measured using a GloMax® 96 Microplate Luminometer (Promega, RRID:SCR_018614). The results are expressed in **Supplementary Fig. 1H** normalized to the cells where negative control LNA is transfected.

Sequence of sense oligo: TCGATGGTGGAGATGCCGGGGA

Sequence of antisense oligo: GGCCTCCCCGGCATCTCCACCA

Codon-Reporter luciferase assays for checking tRNA activity were carried out similarly. In this case the psi-CHECK2 plasmid either has no extra codons (EV) or has 6 test codons (GCG or GCA, that are read by the tRNA being measured) inserted downstream of the initiator methionine in frame with the firefly luciferase. LNA anti-tRF oligonucleotide or a sequence scrambled negative control (NC) were co-transfected with the psi-CHECK2 plasmid and the firefly luciferase levels normalized to renilla luciferase levels. The results are expressed in **Supplementary Fig. 3C** normalized to that in cells containing the EV reporter.

### RNA extraction, qRT-PCR and Northern blotting

For RNA extraction, cells were washed with PBS and lysed directly in TRIzol (Thermo Fisher Scientific,15596026). Total RNA was purified using the Direct-zol™ RNA Miniprep kit (Zymo Research, 50-444-628) according to the manufacturer’s instructions.

For qRT-PCR for EMT markers (mRNA), cDNA was synthesized from total RNA using PrimeScript™ 1st Strand cDNA Synthesis Kit (Takara, 6110A) and applied biosystem thermal cycler (Thermo Fisher Scientific, RRID:SCR_018045). qPCR was performed using Applied Biosystems™ PowerTrack™ SYBR Green Master Mix (A46113) with gene-specific primers for *SNAI2* (Slug), *VIM* (vimentin), *TWIST1*, and *CDH2* (N-cadherin) using the Applied Biosystems StepOnePlus Real-Time PCR System (RRID:SCR_015805). Data were collected using StepOne Software (RRID:SCR_014281). Relative expression was calculated by the ΔΔCt method and normalized to β-actin (*ACTB*).

*ACTB* forward primer: 5′-GCTCCTCCTGAGCGCAAG-3′

*ACTB* reverse primer: 5’-CATCTGCTGGAAGGTGGACA-3′

*CDH2* forward primer: 5’-AGGCTTCTGGTGAAATCGCA-3’

*CDH2* reverse primer: 5’-TGGAAAGCTTCTCACGGCAT-3’

*SNAI2* forward primer: 5’-CATCTTTGGGGCGAGTGAGT-3’

*SNAI2* reverse primer: 5’-ATGGCATGGGGGTCTGAAAG-3’

*TWIST1* forward primer: 5’-GGACAGTGATTCCCAGACGG-3’

*TWIST1* reverse primer: 5’-CCTTTCAGTGGCTGATTGGC-3’

*VIM* forward primer: 5’-ATGCTCAAGAGTGTCAGGGC-3’

*VIM* reverse primer:5’-GGCATGAATGAAACCTGAACCT-3’

For qRT-PCR for small RNAs, total RNA was resolved on Invitrogen™ Novex™ TBE-Urea Gels, 15% (Thermo Fisher Scientific, EC68852BOX) in 1X TBE alongside with Low Range ssRNA Ladder (NEB, N0364S) and microRNA Marker (NEB, N2102S). Gels were stained with SYBR Gold (10,000X), and the region corresponding to the desired tRF size was excised. Gel slices were fragmented using Gel Breaker Tubes (IST Engineering, NC0462125), and RNA was eluted in 0.3 M NaCl overnight at 4 °C. Eluates were cleared using Costar® Spin-X® centrifuge tube filters (0.22 µm) (29442-752) and RNA was precipitated with isopropanol in the presence of GlycoBlue (Thermo Fisher Scientific, AM9516), then washed with 80% ethanol. Reverse transcription and qPCR for tRFs were performed using the Mir-X™ miRNA qRT-PCR TB Green® Kit (Takara, 638314) following the manufacturer’s protocol, together with PowerTrack™ SYBR Green Master Mix (A46113), the kit’s universal reverse primer, and custom specific forward primers. hsa-miR-21-5p was used for normalization. Forward primer sequences were:

hsa-miR-21-5p: 5’-TAGCTTATCAGACTGATGTTGA-3’

tRF-3009a: 5’-ACCCCACTCCTGGTACCA-3’

tRF-3021a: 5’-TCCCCGGCATCTCCACCA-3’

tRF-3030a: 5’-TTCCGGCTCGAAGGACCA-3’

hsa-miR-7-5p: 5’-TGGAAGACTAGTGATTTTGTTGTT-3’

hsa-miR-9-5p: 5’-TCTTTGGTTATCTAGCTGTATGA-3’

For northern blot, the total RNA was separated by denaturing on Invitrogen™ Novex™ TBE-Urea Gels, 15% (Thermo Fisher Scientific, EC68852BOX) in 1X TBE. RNAs were transferred to Amersham Hybond-N+ nylon membrane (Cytiva, RPN303B) using a Bio-Rad Trans-Blot semi-dry transfer apparatus at 200 mA for 1 h in 1X TBE. Membranes were UV-crosslinked and pre-hybridized in ExpressHyb solution (Takara, 636833) for ≥30 min at 42 °C, followed by hybridization in ExpressHyb at 42 °C with biotinylated DNA oligonucleotide probes targeting tRFs and U6 as a loading control. tRF probes were used at 50 pmol/mL, while tRNA and U6 probes were used at 10 pmol/mL:

BIO-Leu-3009: /5BiosG/TGGTACCAGGAGTGGGGT

BIO-Ala-3021: /5BiosG/TGGTGGAGATGCCGGGGATC

BIO-Tyr-3030: /5BiosG/TGGTCCTTCGAGCCGGAATC

U6 probe: /5BiosG/TGCGTGTCATCCTTGCGCAG

Biotin-labeled probes were detected using the Thermo Scientific™ Chemiluminescent Nucleic Acid Detection Module Kit (Thermo Fisher Scientific, 89880) according to the manufacturer’s instructions.

### tRNA-Derived Fragment Profiling in TCGA Lower Grade Glioma Small RNA-seq data and Survival Analysis

We obtained authorization to access raw small RNA sequencing data from the TCGA consortium (RRID:SCR_003193). Raw small RNA-seq data from 512 lower Grade Glioma (LGG) patients were downloaded using the Genomic Data Commons (GDC) Data Portal (RRID:SCR_014514). Read quality was assessed using FastQC (https://www.bioinformatics.babraham.ac.uk/projects/fastqc/, RRID:SCR_014583), followed by 3′ adapter identification and removal using Cutadapt (DOI:10.14806/ej.17.1.200, RRID:SCR_011841). Adapter-trimmed reads were aligned to the human reference genome build hg38 using Bowtie (RRID:SCR_005476) [27] to determine library-specific mapped read counts, which were used for normalization.

Previously curated transfer RNA derived fragments (tRF-5s, tRF-3s, and tRF-1s) from diverse human cell lines were obtained from tRFdb [28]. Each annotated tRF was queried for presence and abundance across all libraries. To account for differences in sequencing depth, raw counts were normalized to reads per million mapped reads (RPM).

Clinical metadata, including overall survival and vital status, were retrieved from the TCGA portal using the Genomic Data Commons [29] R package (https://bioconductor.org/packages/GenomicDataCommons). Patients with incomplete survival information were excluded, resulting in 465 evaluable cases. For each tRF, patients were ranked by normalized expression and stratified into tertiles (top, middle, bottom) to assess potential dose-dependent associations while maintaining statistical power.

Associations between tRF expression and overall survival were assessed using univariate Cox proportional hazards regression, comparing patients in the top-third versus the bottom-third expression groups for each tRF. Kaplan–Meier survival curves were generated, and significance was assessed using the log-rank test. All survival analyses and visualizations were performed in R (R Project for Statistical Computing, RRID:SCR_001905) using the survival package (RRID:SCR_021137) and survminer package (RRID:SCR_021094).

### RNA-seq library preparation and data analysis

#### RNA-seq Alignment and Quantification

8 h: Total RNA from knockdown (KD) and negative control (NC) cells was submitted to Plasmidsaurus for 3’-RNA-seq. First-strand cDNA was generated by reverse transcription using an oligo(dT) primer to selectively capture polyadenylated transcripts, followed by second-strand synthesis, tagmentation, and library preparation. Libraries were quantified by Qubit and assessed on an Agilent TapeStation (library kit details are proprietary to Plasmidsaurus). Libraries were sequenced on an Illumina NovaSeq X, generating ∼20 million raw single-end reads per sample. Reads were processed through UMI-based deduplication to yield ∼10 million deduplicated 3’-end reads per sample. Bioinformatic processing was performed using a standard RNA-seq pipeline: reads were demultiplexed (BCL Convert), filtered/trimmed using fastp (version 0.23.4, RRID:SCR_016962) [30], aligned to the human reference genome (GRCh38, primary assembly) using HISAT2 (version 2.2.1, RRID:SCR_015530) [31], sorted using samtools (version 1.12, RRID:SCR_002105) [32], and UMI-deduplicated using UMIcollapse (version 1.1.0). Gene-level counts were generated with featureCounts (Subread package, version 2.1.1, RRID:SCR_012919) (Dolgalev I. msigdbr: MSigDB Gene Sets for Multiple Organisms in a Tidy Data Format. R package version 7.5.1).

48 hr: Sequence reads were filtered/trimmed using fastp (version 0.23.4), and aligned to the human reference genome (GRCh38, primary assembly) using HISAT2 (version 2.2.1) [30]. The genome index was constructed based on the GRCh38 primary assembly. Mapping was performed with reverse-strand-specific settings, and all other alignment parameters were set to default values. The resulting SAM files were converted to BAM format and sorted by coordinate using SAMtools (version 1.12) [31]. Gene expression levels were quantified using featureCounts (version 2.1.1, Subread package) [32]. Reads were assigned to genes based on the MANE v1.4 annotation (Ensembl genomic GTF). Multi-mapping reads were excluded, and counting was performed in a strand-specific manner matching the alignment protocol.

#### Differential Gene Expression Analysis

Differential gene expression analysis was performed using the R package edgeR (version 3.34.0, RRID:SCR_012802) [33]. Genes with low expression levels across all samples were removed using the filterByExpr function. The filtered count matrix was normalized using the Trimmed Mean of M-values (TMM) method. Biological variability was estimated using estimateDisp, and a generalized linear model (GLM) was fitted using glmQLFit. Differential expression was assessed using a quasi-likelihood F-test. Genes satisfying an FDR threshold of <0.05 were considered differentially expressed.

#### Visualization of Differential Expression

To visualize global transcriptional changes and compare results between contrasts, the following plots were generated:

#### Volcano Plot

Volcano plots were generated using ggplot2 (version 3.4.4, RRID:SCR_014601) [34]. The x-axis represents logFC and the y-axis represents -log10(FDR). Genes were color-coded based on regulation status: upregulated genes (logFC > 1.0, FDR < 0.05) are shown in orange, and downregulated genes (logFC < -1.0, FDR < 0.05) are shown in purple. Labels were added to the most significant genes.

#### Venn Diagrams

Venn diagrams were generated using ggvenn (version 0.1.10) to identify common and unique differentially expressed genes (DEGs) between comparisons. Separate diagrams were generated for upregulated and downregulated genes.

#### Gene Set Enrichment Analysis (GSEA)

GSEA was performed to identify biological pathways associated with changes in gene expression. A ranked gene list was generated using all detected genes, with the ranking metric calculated as:

Rank = sign(log2FC) × -log10(P-value)

This metric incorporates both the direction of change (sign of logFC) and the statistical significance (P-value). Ensembl gene IDs were mapped to Entrez IDs using the bitr function of clusterProfiler (RRID:SCR_016884) [35]. Duplicate Entrez IDs arising from one-to-many mappings were removed, and entries with the highest ranking metric were retained. GSEA was performed using ClusterProfiler and the msigdbr package (Dolgalev I. msigdbr: MSigDB Gene Sets for Multiple Organisms in a Tidy Data Format. R package version 7.5.1.), with 1,000 permutations. The Reactome pathway database (RRID:SCR_003485) and MSigDB Hallmark (H) gene sets (RRID:SCR_016863) were queried. Top 20 significant pathways (adjusted P-value < 0.05) were visualized in a bar graph, where the x-axis represents the normalized enrichment score (NES), and bars were color-coded based on Benjamini–Hochberg adjusted p-values.

### Statistical Analysis

Statistical analyses were performed using GraphPad Prism version 8.0.2 (GraphPad Software, RRID:SCR_002798). Data are presented as mean ± SD unless otherwise indicated. Two-group comparisons were analyzed using unpaired two-tailed t-tests. Comparisons among multiple groups were analyzed using one-way ANOVA, or two-way ANOVA for time-course experiments, followed by the appropriate multiple-comparisons tests as indicated in the figure legends. Where applicable, multiple testing was controlled using the Benjamini–Krieger–Yekutieli false discovery rate approach. P < 0.05 was considered statistically significant. The number and type of replicates are indicated in the figure legends.

### Data Availability

RNA-seq data have been submitted to the Gene Expression Omnibus (GEO), and the accession number will be provided once available.

## Results

### High expression of tRF-3s is associated with poor prognosis in LGG patients

To identify tRFs that could be important for cancer biology, we analyzed the short RNA-seq data of TCGA cancers to determine whether levels of any of the tRF-3s correlate with a change in outcome. This survival analysis was performed as a discovery and prioritization screen to identify tRFs warranting functional investigation. Patients were stratified into top-versus-bottom tertile of tRF expression.

A strong signal was detected in lower grade gliomas (LGG) where levels of three tRF-3s correlated with patient survival **(Fig. 1A)**. High levels of tRF-3009a, tRF-3021a and tRF-3030a (and not other tRF-3s) are associated with poor outcome in overall survival **(Fig. 1B-D)**. These tRFs are 18 nucleotides tRNA fragments processed from cleavage in TψC-loop of mature tRNA and extend to the CCA end **(Fig. 1E)** and have been categorized as tRF-3a (1, 2). tRF-3009a, tRF-3021a and tRF-3030a are derived from tRNA-Leu-TAA, tRNA-Ala-TGC and tRNA-Tyr-GTA, respectively. None of these tRFs have been investigated before in the context of glioma biology.

### Experimental manipulation of tRF levels in cell lines

The tRFs were knocked down (KD) in cells by mixmer locked nucleic acid (LNA) antisense oligonucleotides while their overexpression (OE) was achieved by transfection of synthetic mimics of the tRFs. The specificity of KD and OE was confirmed by qPCR **(Supplementary Fig. 1A-F)**. In addition, LNA KD did not decrease parental tRNA expression, though in two cases there seems to be an increase in the tRNA levels **(Supplementary Fig. 1G)**. These data demonstrate that our LNA and mimic reagents efficiently and specifically modulate tRF-3 levels in U251 cells.

We know from our previous studies that tRF-3s repress luciferase and endogenous cellular genes that contain a sequence complementary to the tRFs [12, 13]. Thus, as an orthogonal functional validation of tRF-3021a knockdown, we performed a dual luciferase reporter assay in which the reporter harbors a sequence complementary to tRF-3021a. The level of repression of this reporter will be indicative of the level of functional tRF-3021a in the cell. LNA-mediated knockdown of tRF-3021a de-repressed luciferase activity specifically **(Supplementary Fig. 1H)** confirming efficient depletion of endogenous tRF-3021a. Consistent with the reporter being sensitive to the level of the tRF-3021a, transfection of the tRF-mimic repressed the luciferase further. This orthogonal validation of the knockdown was done only with tRF-3021a because of the profound effects of knockdown of this tRF that we will report below.

### tRF-3 fragments enhance cell invasion and migration of GBM cells

Given the correlation between high tRF-3 expression and poor prognosis, we wondered whether these tRFs promote invasion and migration of cancer cells. Since there are no cell lines available that are derived from LGGs, we initiated our studies with established cell lines obtained from glioblastoma cells. The data shown here focus on U251 cells, similar effects were also observed in U2OS and DU145 cells, suggesting that the pro-migratory and pro-invasive effects of these tRFs are not restricted to GBM cells (data not shown).

Invasion and migration of U251 cells were measured by Transwell assays after the KD or OE of the relevant tRFs. In all assays, values were normalized to the corresponding Negative Control (NC: LNA or mimic). KD of tRF-3009a, tRF-3021a and tRF-3030a reduced invasion and migration in 18-24 hours of U251 cells after seeding into the Transwell inserts, with the strongest effect observed for tRF-3021a **(Fig. 2A–C)**. KD of tRF-3009a, tRF-3021a and tRF-3030a reduced invasion to 50.96%, 0.59% and 40.64%, respectively **(Fig. 2B)**. KD of tRF-3009a and tRF-3021a also markedly reduced migration to 42.21% and 0.47%, respectively, whereas tRF-3030a KD produced a more modest and variable reduction to 65.00% **(Fig. 2C)**. Conversely, OE of tRF-3009a and tRF-3021a enhanced invasion to 191.4% and 236.6%, respectively, while tRF-3030a OE showed a smaller increase to 168.5% **(Fig. 2D, E)**. OE of tRF-3009a and tRF-3021a similarly increased migration to 179.2% and 193.9%, respectively, whereas tRF-3030a OE produced a modest change to 120.7% **(Fig. 2D, F)**. To demonstrate that the reduction in invasion and migration reflects a specific pro-invasive function of tRF-3021a rather than a general collapse in cell fitness, we performed Transwell assays at an earlier 12-hour time point, prior to the onset of the significant apoptosis that will be reported below. We transfected the cells with 30nM tRF-3021a LNA for KD and 30nM tRF-3021a LNA and 100nM tRF-3021a mimic for rescue. tRF-3021a KD reduced both invasion and migration at 12 hours, and co-transfection of a tRF-3021a mimic rescued both phenotypes **(Supplementary Fig. 1I-K)**.

To see if changes in invasion and migration are associated with a shift to the mesenchymal/invasive-state, we measured markers of the invasive state in U251 cells. qRT-PCR analysis showed that anti-tRF-3021a reduced the expression of invasive-state markers, including *CDH2*, *SNAI2*, *VIM* and *TWIST1* mRNA levels **(Fig. 2G)**. To determine whether the shift away from the mesenchymal state after tRF-3021a knockdown occurred at the protein level, we examined N-cadherin, SLUG and TWIST1 by immunoblotting. Consistent with the qRT-PCR data, tRF-3021a knockdown reduced the protein abundance of N-cadherin and TWIST1 and, most robustly, SLUG **(Fig. 2H**). SLUG was likewise reduced upon tRF-3009a or tRF-3030a knockdown, consistent with shared invasion/migration effects. These results support that tRF-3021a promotes a pro-invasive mesenchymal program in GBM cells. Together, these data identify tRF-3s as novel regulators that are important for migration and invasion of GBM cells.

### tRF-3021a was uniquely required for cell proliferation and suppression of apoptosis

Cell proliferation measured by MTT showed that tRF-3021a KD slowed proliferation in U251 cells, with significantly reduced viability at 48 h and 72 h, whereas tRF-3009a KD and tRF-3030a KD had minimal or no effect at any time point **(Fig. 2I)**. Consistent with this, tRF-3021a KD also decreased MTT signal in normal human astrocytes and U87 glioma cells **(Supplementary Fig. 1L, M)**, indicating that tRF-3021a promotes proliferation in both non-neoplastic and malignant glial cells. Because the MTT assay measures metabolic activity rather than cell number directly, and because tRF-3021a KD alters both protein synthesis and oxidative phosphorylation related gene signatures which we will discuss later, we also directly counted the cells at different time points. Cell counts at 24, 48, and 72 hours confirmed a significant reduction in cell number in tRF-3021a KD compared with NC cells by 24 hours **(Fig. 2J),** providing direct evidence of reduced proliferation independent of metabolic readout.

To further explore the role of tRF-3021a in cell viability and survival, we asked whether the loss of tRF-3021a cause any cell-cycle block in U251 cells by analyzing the DNA content 48 hours post transfection with NC LNA or tRF-3021a LNA. tRF-3021a KD caused an accumulation of a sub-G1 population, with little effect on G1, S or G2/M, plotted after subtracting the sub-G1 population **(Fig. 3A–D)**. The most striking change was in the sub-G1 fraction, which increased from 1.3% ± 0.8% to 47.9% ± 6.1%, indicating extensive apoptosis.

**Figure 3.**
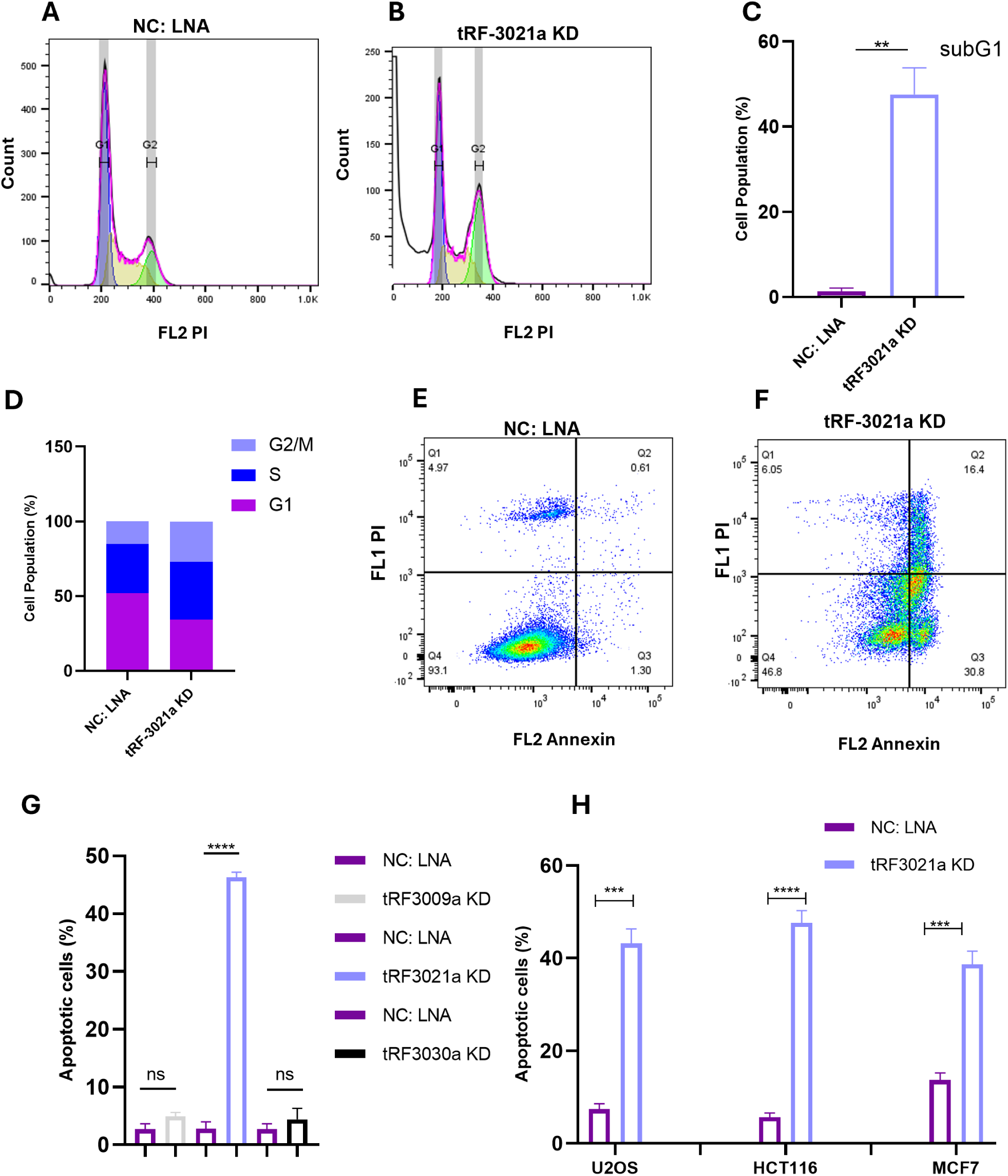
tRF-3021a knockdown induces apoptosis in GBM cells and non-GBM cells. (A–B) Representative propidium iodide (PI) DNA-content histograms for cell-cycle profiling of U251 cells 48 h after transfection with negative-control (NC) LNA (A) or tRF-3021a LNA (B). (C–D) Quantification of the sub-G1 population (C) and cell-cycle distribution (D) in U251 cells after tRF-3021a knockdown (KD) compared with NC LNA. (E–F) Representative Annexin V/PI flow-cytometry plots of U251 cells 48 h after transfection with NC LNA (E) or tRF-3021a LNA (F). Quadrants indicate live cells, early apoptotic cells, late apoptotic/necrotic cells, and PI-only populations. (G) Quantification of Annexin V-positive cells, including early and late apoptotic cells, in U251 cells after KD of tRF-3009a, tRF-3021a, or tRF-3030a compared with the corresponding NC LNA. (H) Annexin V-positive cells after tRF-3021a KD compared with NC LNA in U2OS, HCT116, and MCF7 cells. Data are presented as mean ± SD from independent biological replicates unless otherwise indicated. Statistical significance was determined by unpaired two-tailed t-test for the sub-G1 comparison and by multiple unpaired two-tailed t-tests with Benjamini–Krieger–Yekutieli false-discovery-rate correction for Annexin V comparisons. ns, not significant, *P < 0.05, **P < 0.01, ***P < 0.001, ****P < 0.0001.

Because of the striking increase in sub-G1 cells, we quantified apoptosis by Annexin V/PI staining 48 hours post transfection. tRF-3021a KD increased the percentage of Annexin V–positive U251 cells from 2.8% in NC LNA to 46.3% in tRF-3021a LNA, while tRF-3009a and tRF-3030a KD did not induce apoptosis **(Fig. 3E-G)**. The same effect was observed in non-GBM cells: U2OS (osteosarcoma), HCT116 (colon cancer) and MCF7 (breast cancer) **(Fig. 3H)** and in normal astrocytes and U87 GBM cells **(Supplementary Fig. 1N)**.

Together, these data indicate that depletion of tRF-3021a promotes apoptosis in GBM and non-GBM cells. Based on the very strong phenotype of tRF-3021a on migration, invasion, proliferation and apoptosis, we focused subsequent molecular studies on tRF-3021a.

### tRF-3021a knockdown inhibits global protein synthesis

Considering tRF-3021a requirement for cell proliferation, survival and protection from apoptosis, we wondered whether tRF-3021a somehow contributes to the maintenance of normal protein synthesis, which is required for cell growth and survival. To assess global translation rates, we performed puromycin-labeling assays following knockdown of tRF-3021a. tRF-3021a KD led to a strong reduction in puromycin incorporation in U251 cells at 48 h post knockdown, indicating decreased global protein synthesis. A similar reduction in global protein synthesis was also observed in multiple glioma and non-glioma cell lines, including normal astrocytes, U87, A172, U2OS (osteosarcoma), HCT116 (colon cancer), and MCF7 (breast cancer) **(Fig. 4A)**. In contrast, knockdown of tRF-3009a or tRF-3030a had minimal impact on translation, highlighting the unique role of tRF-3021a in sustaining protein synthesis **(Fig. 4B)**. To confirm these findings with an orthogonal metabolic labeling approach, we performed L-homopropargylglycine (HPG) incorporation assays, which detect nascent protein synthesis independently of puromycin. HPG incorporation was similarly reduced in tRF-3021a-depleted U251 cells compared with NC-treated cells showing after 24 hours post-transfection **(Fig. 4C)**, providing independent orthogonal confirmation of the translational inhibition phenotype. We also demonstrated the block to protein synthesis with an independent gapmer LNA targeting the full tRF-3021a sequence. Consistent with the mixmer anti-tRF-3021a results, gapmer-mediated knockdown decreased global protein synthesis, and qRT-PCR confirmed efficient depletion of tRF-3021a **(Fig. 4D, E)**, providing a second orthogonal reagent that confirms that this profound block in protein synthesis is due to tRF-3021a knockdown and not due to some contaminants present in the mixmer anti-tRF-3021a.

**Figure 4.**
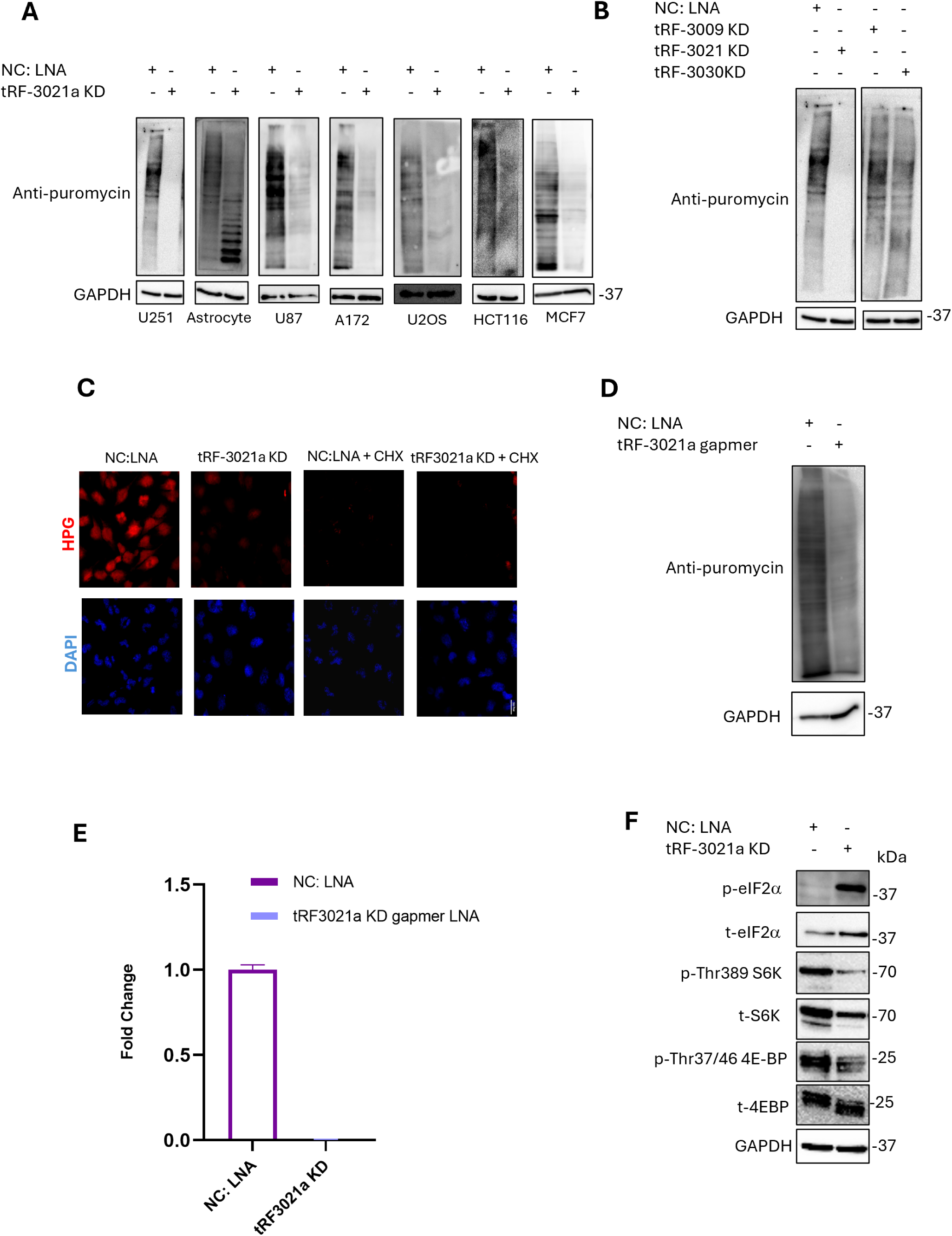
tRF-3021a knockdown inhibits global protein synthesis. (A) Puromycin incorporation assay in the indicated cell lines after LNA-mediated knockdown (KD) of tRF-3021a compared with negative-control (NC) LNA. Cells were incubated with puromycin, and lysates were immunoblotted with anti-puromycin to assess global protein synthesis. GAPDH was used as a loading control. (B) Puromycin incorporation assay in U251 cells after KD of tRF-3009a, tRF-3021a, or tRF-3030a compared with NC LNA. GAPDH was used as a loading control. (C) HPG incorporation assay in U251 cells after tRF-3021a KD compared with NC LNA 24 h after transfection. Cycloheximide-treated cells were used as a negative control for translation. DAPI was used to stain nuclei. (D) Puromycin incorporation assay in U251 cells after gapmer-mediated KD of tRF-3021a compared with NC LNA, showing reduced global protein synthesis using an independent knockdown strategy. GAPDH was used as a loading control. (E) qRT-PCR validation of tRF-3021a reduction in U251 cells after transfection with tRF-3021a gapmer LNA compared with NC LNA. Expression was normalized to NC LNA. (F) Immunoblot analysis of translation-regulatory signaling in U251 cells after tRF-3021a KD compared with NC LNA. Total and phosphorylated eIF2α, p70S6K, and 4E-BP1 were assessed. GAPDH was used as a loading control. Panel E is shown as mean ± SD from technical qPCR replicates.

To gain insight into the pathways driving the reduced protein synthesis we examined phosphorylation of several translational regulators at different time points upon tRF-3021a KD. Immunoblotting showed decreased phosphorylation of substrates of the mTOR pathway, including p70S6 kinase (Thr389) and 4EBP1 (Thr37/46), indicating that mTORC1 signaling decreased after 48 h of tRF-3021a knockdown (**Fig. 4F)**. A slight increase of total and a marked increase of p-eIF2α at 48 hours after KD suggest activation of the integrated stress response (ISR) **(Fig. 4F)**.

Together, these findings demonstrate that tRF-3021a is required for maintaining basal levels of protein synthesis.

### Translation inhibition precedes and is independent of apoptosis

To determine whether protein synthesis inhibition is downstream or upstream of apoptosis, we investigated Annexin V–positive cells and puromycin incorporation at different time points after tRF-3021a knockdown (2–48 h). Apoptosis began to increase as early as 12 h and became prominent at 48 h **(Fig. 5A)**, correlating with increased cleaved PARP **(Fig. 5B)**. In contrast, protein synthesis showed a decrease starting as early as 8 h post knockdown before significant apoptosis **(Fig. 5C, representative blot shown in Supplementary Fig. 2A)**. Consistent with this, the decrease in p-S6K and p-4EBP is seen as early as at 4 h post knockdown **(Fig. 5D)**. Note that in contrast, the activation of the integrated stress response (as seen by pEIF2α) is not significant even at 12 h post-transfection, suggesting that activation of ISR is unlikely to be the first cause of the protein synthesis inhibition (**Fig. 5E**).

**Figure 5.**
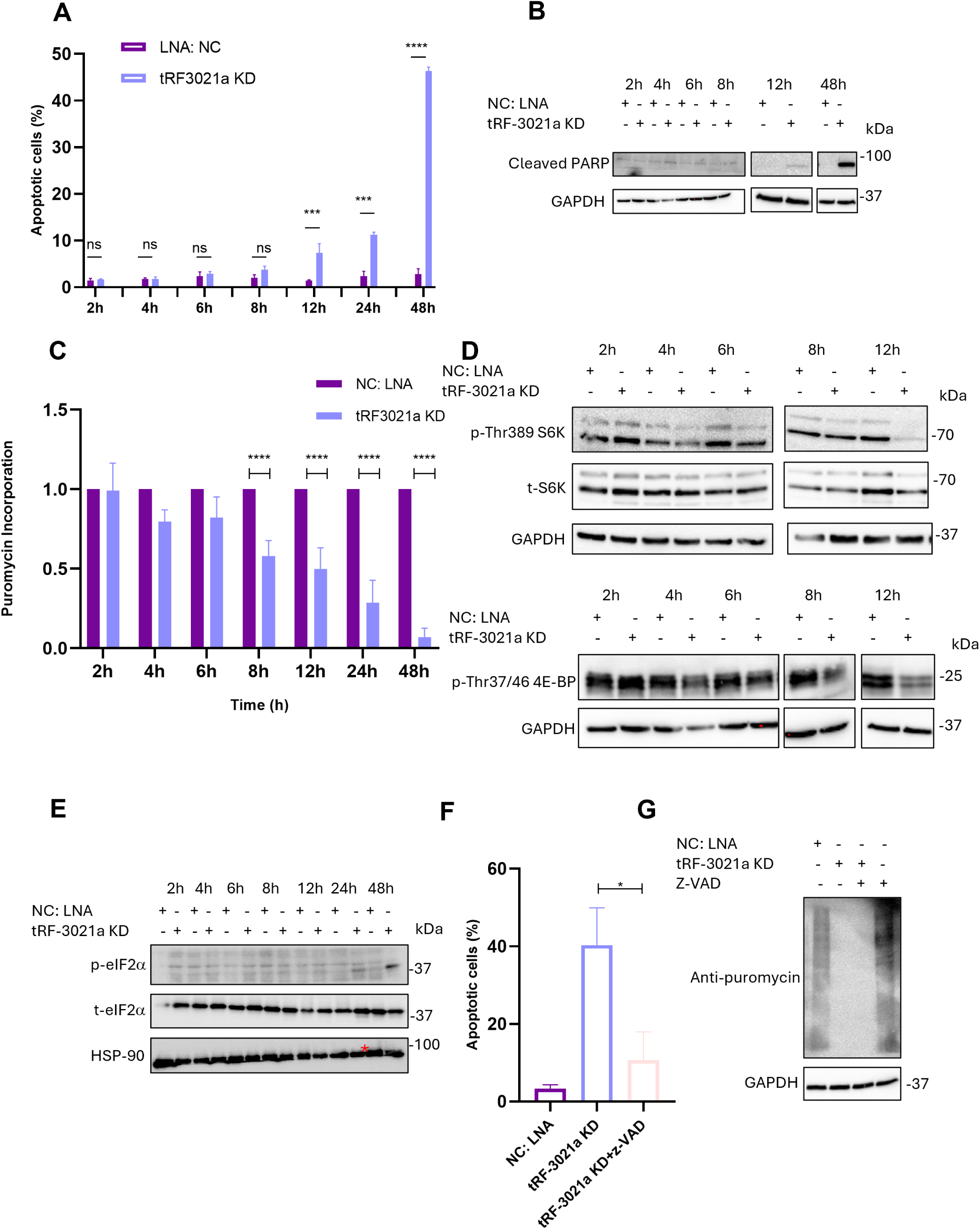
Translation inhibition precedes and is independent of apoptosis. (A) Time-course analysis of Annexin V-positive apoptotic cells in U251 cells after LNA-mediated knockdown (KD) of tRF-3021a compared with negative-control (NC) LNA at the indicated time points. (B) Immunoblot analysis of cleaved PARP in U251 cells after tRF-3021a KD compared with NC LNA at the indicated time points. GAPDH was used as a loading control. (C) Quantification of puromycin incorporation in U251 cells after tRF-3021a KD compared with NC LNA at the indicated time points. Values were normalized to NC LNA at each time point. (D) Immunoblot analysis of p70S6K and 4E-BP1 phosphorylation in U251 cells after tRF-3021a KD compared with NC LNA at the indicated time points. GAPDH was used as a loading control. (E) Immunoblot analysis of total and phosphorylated eIF2α in U251 cells after tRF-3021a KD compared with NC LNA at the indicated time points. HSP90 was used as a loading control. (F) Quantification of Annexin V-positive apoptotic cells in U251 cells after NC LNA, tRF-3021a KD, or tRF-3021a KD with z-VAD-fmk treatment. (G) Puromycin incorporation assay in U251 cells after NC LNA, tRF-3021a KD, NC LNA with z-VAD-fmk, or tRF-3021a KD with z-VAD-fmk treatment. GAPDH was used as a loading control. Data are presented as mean ± SD from independent biological replicates unless otherwise indicated. Statistical significance for the time-course experiments in panels A and C was determined by two-way ANOVA with multiple-comparisons testing comparing tRF-3021a KD with NC LNA at each time point. Statistical significance for panel F was determined by one-way ANOVA with multiple-comparisons testing. Immunoblots are representative experiments unless otherwise indicated. ns, not significant, *P < 0.05, **P < 0.01, ***P < 0.001, ****P < 0.0001.

Because protein synthesis inhibition preceded the earliest signs or apoptosis, we suspected that that is the primary defect and is not the result of apoptotic pathway induction. To test this, we treated cells with the pan-caspase inhibitor z-VAD-fmk which substantially reduced Annexin V-positive cells in tRF-3021a KD samples **(Fig. 5F)** but failed to restore puromycin incorporation **(Fig. 5G)**. These findings indicate that protein synthesis inhibition happens earlier than apoptosis (8 h vs. 12 h) and is independent of apoptosis, as z-VAD could not restore protein synthesis.

### Apoptosis induction and protein synthesis inhibition upon tRF-3021a KD can be restored by introducing tRF-3021a mimic

To confirm the specificity of these effects and verify that the observed phenotypes are due to loss of tRF-3021a, we performed rescue experiments. We introduced synthetic tRF-3021a mimic with the LNA. tRF-3021a mimic could rescue the phenotypes caused by the LNA mediated knockdown. Annexin V/PI analysis showed that tRF-3021a mimic alone did not alter apoptosis, while co-transfection of the tRF-3021a mimic with tRF-3021a LNA significantly reduced the proportion of apoptotic cells compared to knockdown alone **(Fig. 6A)**. Similarly, puromycin incorporation was restored upon tRF-3021a mimic rescue **(Fig. 6B)**. To further rule out nonspecific effects related to depletion of a highly abundant small RNA (given that tRF-3021a is more abundant in U251 cells than tRF-3009a and tRF-3030a), we inhibited two abundant U251 miRNAs (miR-7-5p and miR-9-5p) using the same LNA-based approach. Knockdown of miR-7-5p and miR-9-5p was confirmed by qRT-PCR **(Supplementary Fig. 2B, C)**. In contrast to tRF-3021a knockdown, inhibition of these miRNAs did not reduce puromycin incorporation **(Fig. 6C)**, indicating that global translation inhibition is not simply an abundance-dependent outcome of depleting a highly expressed small RNA. Collectively, these data support that the observed apoptosis induction and protein synthesis inhibition are specific to tRF-3021a knockdown rather than off-target effects of the LNA.

**Figure 6.**
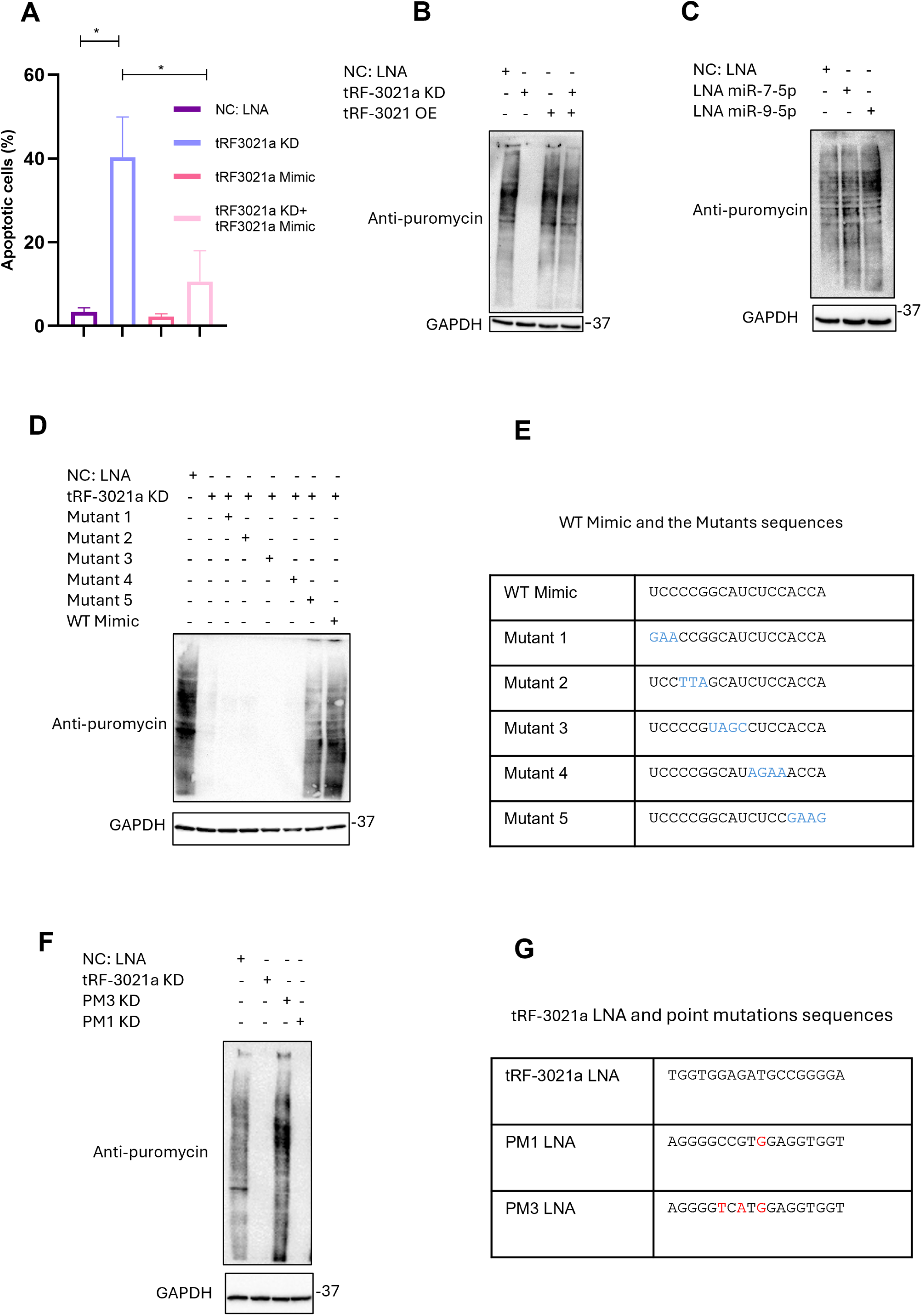
Apoptosis induction and protein synthesis inhibition upon tRF-3021a KD can be restored by introducing tRF-3021a mimic. (A) Quantification of Annexin V-positive apoptotic cells in U251 cells transfected with negative-control (NC) LNA, tRF-3021a LNA, tRF-3021a mimic, or tRF-3021a LNA together with tRF-3021a mimic. (B) Puromycin incorporation assay in U251 cells transfected with NC LNA, tRF-3021a LNA, tRF-3021a mimic, or tRF-3021a LNA together with tRF-3021a mimic. GAPDH was used as a loading control. (C) Puromycin incorporation assay in U251 cells after LNA-mediated knockdown (KD) of miR-7-5p or miR-9-5p compared with NC LNA. GAPDH was used as a loading control. (D) Puromycin incorporation assay in U251 cells transfected with NC LNA, tRF-3021a LNA, or tRF-3021a LNA together with WT or mutant tRF-3021a mimics. GAPDH was used as a loading control. (E) Sequences of the WT tRF-3021a mimic and mutant tRF-3021a mimics used in panel D. Mutated nucleotides are shown in blue. (F) Puromycin incorporation assay in U251 cells transfected with NC LNA, tRF-3021a LNA, or point-mutated tRF-3021a LNAs. GAPDH was used as a loading control. (G) Sequences of tRF-3021a LNA and point-mutated tRF-3021a LNAs used in panel F. Mutated nucleotides are shown in red. Data are presented as mean ± SD from independent biological replicates unless otherwise indicated. Statistical significance was determined by one-way ANOVA with multiple-comparisons testing for the apoptosis rescue experiment. ns, not significant, *P < 0.05, **P < 0.01, ***P < 0.001, ****P < 0.0001.

To test the sequence-specificity of the mimic rescue, we performed rescue experiments with a panel of tRF-3021a mimics carrying mutations at different positions. Mutations in the first 14 residues of the tRF-3021a abrogated the ability of the mimic to rescue protein synthesis. Only Mutant 5 with mutations in the last three bases at the 3′ end, could still rescue puromycin incorporation **(Fig. 6F, G)**. Tm calculations indicated that all the mutants can efficiently anneal with and titrate the anti-tRF-3021a LNA at the cell culture temperatures. These results demonstrate that the rescue is sequence-dependent and reflects functional restoration of tRF-3021a activity, not non-specific LNA neutralization by the tRF mimics simply annealing to the LNA.

We also tested the sequence-specificity of the LNA in protein synthesis inhibition by designing two mutant LNAs, targeting tRF-3021a: PM1 carries a single base mutation while PM3, carries three base mutations relative to the original LNA targeting tRF-3021a. PM3 failed to inhibit translation, whereas PM1 retained the ability to reduce global protein synthesis **(Fig. 6H, I)**. PM1 can decrease tRF-3021a to the same extent as tRF-3021a LNA, while PM3 shows less decrease in tRF-3021a levels **(Supplementary Fig. 2D)**. These results suggest that the residual tRF-3021a after PM3-mediated knockdown is sufficient to sustain protein synthesis and that the inhibition of protein synthesis by the LNA is dependent on its ability to titrate down the tRF-3021a.

Together, the mutations in the tRF-3021a mimic and the anti-tRF-3021a LNA establish that the translational inhibition following tRF-3021a knockdown is attributable to the specific loss of tRF-3021a activity.

### tRF-3021a-mediated translation inhibition is not caused by disruption or depletion of the parental tRNA-Ala

Previous experiments by us and others have suggested that an anti-tRF does not target the parental tRNA [26, 36] most likely because the secondary structure of the tRNA makes it inaccessible to the anti-tRF. Indeed, Northern blots did not show any decrease of tRNA-Ala upon transfection of anti-tRF-3021a (**Supplementary Fig. 1G**). Nevertheless, we wanted additional evidence that the inhibition of protein synthesis is not because the anti-tRF-3021a is annealing to tRNA-Ala-TGC, the parent to tRF-3021a. First, we designed two additional LNAs targeting other regions of the parental tRNA-Ala including, a 5′- LNA targeting the 5′ end of the tRNA-Ala and an anti-codon LNA targeting the anticodon loop of tRNA-Ala **(Supplementary Fig. 3A)**. Neither the anti-5′-LNA nor the anti-codon LNA reduced global protein synthesis as assessed by puromycin incorporation (Supplementary Fig. 3B).

Second, we performed a codon reporter assay in which efficient translation of the reporter depends on functional tRNA-Ala-TGC to decode GCA codons and another that requires tRNA-Ala-CGC to decode GCG codons as control. This is achieved by inserting a trail of six codon in the 5’ end of the ORF of the firefly luciferase. If the relevant tRNA is depleted, the efficiency of the translation of the codon reporter will be selectively decreased compared to the negative control reporter that has WT firefly luciferase sequence without the insertion of the six codons (EV). tRF-3021a knockdown did not impair GCA-codon or GCG-codon reporter activity **(Supplementary Fig. 3C)**, confirming that global protein synthesis inhibition is not a consequence of disrupting tRNA-Ala activity. Taken together, these experiments demonstrate that the translational phenotype is specifically attributable to loss of the tRF-3021a fragment itself.

### Transcriptomic changes following tRF-3021a knockdown

To identify genes associated with tRF-3021a–mediated invasion, migration, and apoptosis in GBM, U251 cells were transfected with an LNA targeting tRF-3021a to achieve knockdown, and RNA-seq was performed at 8 and 48 h after transfection, with three biological replicates per condition at each time point. Each time point included three NC and three tRF-3021a KD biological replicates. Differential expression analysis revealed that tRF-3021a depletion produced a distinct gene expression (DEG) signature at both time points compared with control cells **(Fig. 7A–F)**. Using FDR < 0.05 and |log2FC| ≥ 1 thresholds, we identified 969 upregulated and 2,520 downregulated genes at 8 h **(Fig. 7A)**, and 1,287 upregulated and 1,245 downregulated genes at 48 h **(Fig. 7B) (Supplemental Table 1)**.

**Figure 7.**
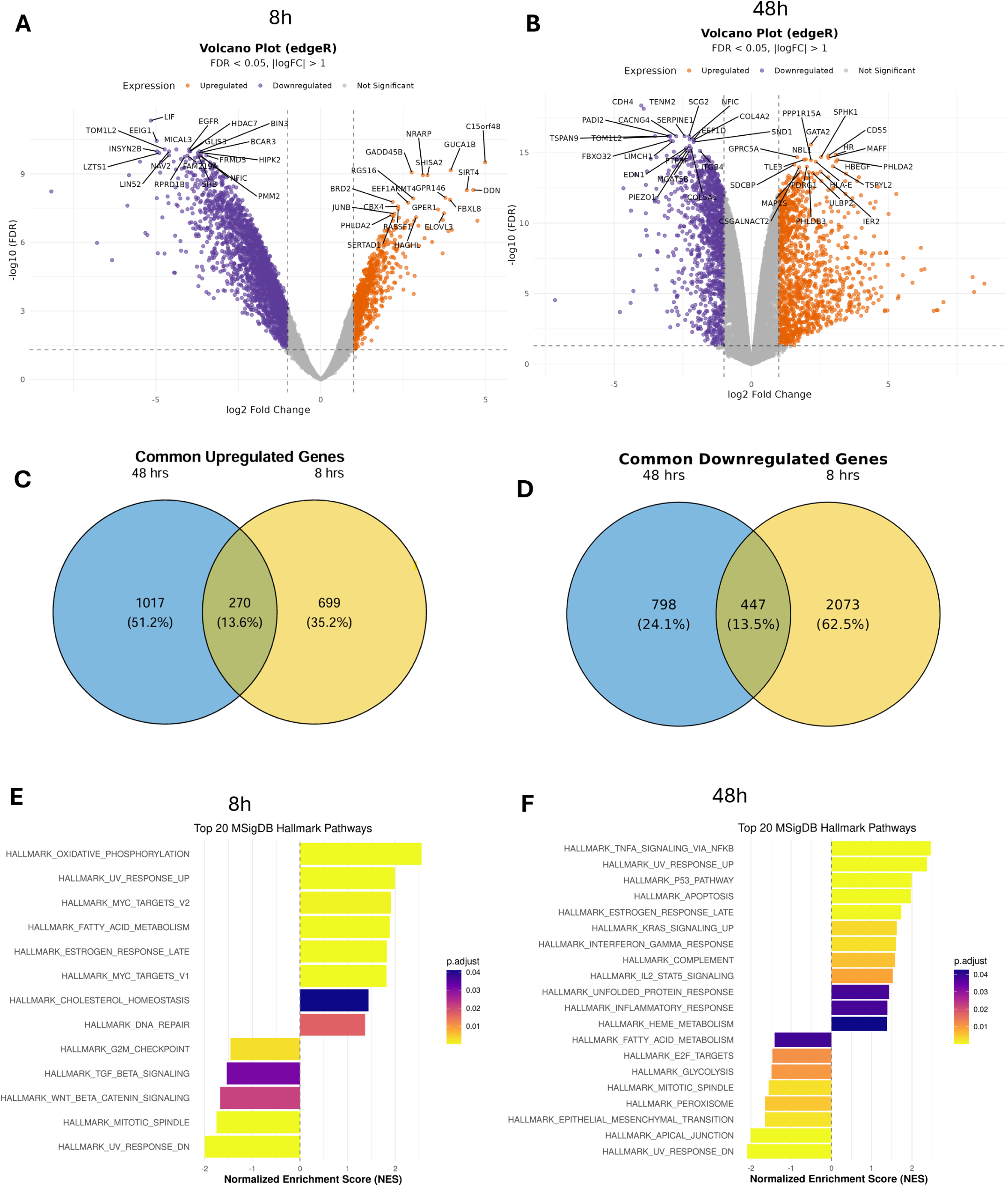
Transcriptomic changes following tRF-3021a knockdown. (A–B) Volcano plots showing differentially expressed genes in U251 cells after LNA-mediated knockdown (KD) of tRF-3021a compared with negative-control (NC) LNA at 8 h (A) and 48 h (B). Upregulated genes are shown in orange, downregulated genes are shown in purple, and non-significant genes are shown in gray. Differentially expressed genes were defined using FDR < 0.05 and |log2 fold change| > 1. (C–D) Venn diagrams showing the overlap of upregulated genes (C) and downregulated genes (D) between the 8 h and 48 h tRF-3021a KD RNA-seq datasets. (E–F) MSigDB Hallmark pathway enrichment analysis of genes altered after tRF-3021a KD at 8 h (E) and 48 h (F). Normalized enrichment scores are shown, and adjusted P values are indicated by color. Differential gene expression analysis was performed using edgeR with FDR correction for multiple testing. Hallmark pathway enrichment was analyzed using adjusted P values.

We next compared DEGs between time points to define common DEGs versus unique DEGs for each time point. Among upregulated genes, 270 were common in both 8 and 48 hours, while among downregulated genes, 447 were shared **(Fig. 7C-D)**. This comparison indicates that tRF-3021a depletion drives substantial differential gene-expression and a transition from early transcriptomic changes when translation inhibition begins to appear to later changes when apoptosis is more pronounced.

To study biological pathways associated with these time-dependent signatures, we performed gene set enrichment analysis (GSEA) using MSigDB Hallmark gene sets. At 8 h, the most positively enriched pathways included oxidative phosphorylation (OXPHOS) and UV_RESPONSE_UP, whereas negatively enriched pathways included multiple cell cycle–associated signatures, as well as TGF-β and WNT signaling programs **(Fig. 7E)**. In contrast, at 48 h, the most positively enriched pathways included TNFα signaling via NF-κB, p53 pathway, and apoptosis **(Fig. 7F)**. TNFα signaling via NF-κB signaling was among the most enriched programs at both time points **(Supplementary Fig. 4A)**. Reactome GSEA further showed that at 8 h post-knockdown genes related to eukaryotic translation initiation/elongation and response to amino acid deprivation were strongly enriched **(Supplementary Fig. 4B)**, whereas at 48 h, we observed enrichment of necrosis and cell death–related pathways **(Supplementary Fig. 4C)**. At 8 h, gene ontology analysis by molecular function (GO_MF) revealed a strong enrichment among up-regulated genes of those involved in ribosomal function and mitochondrial respiratory and oxidative phosphorylation programs, including electron transport chain and ATP synthesis–coupled respiration related genes **(Supplementary Fig. 4D)**.

Collectively, these results show that early after knockdown there is a disruption of translational homeostasis, oxidative phosphorylation, DNA damage response and cell-cycle progression pathways, but later the gene expression program is suggestive of terminal apoptosis and necrosis with activation of p53 mediated signaling.

tRF-3s can exert microRNA-like silencing activity, where they repress endogenous cellular genes carrying microRNA-like target sequences following standard seed-sequence rules [13]. Therefore, we examined the RNA-seq data for genes de-repressed by the anti-tRF-3021a by such seed-sequence rules that might explain the early inhibition of protein synthesis. No such genes were found (data not shown), suggesting that the mechanism of inhibition of protein synthesis by anti-tRF-3021a cannot yet be attributed to the microRNA like functions of tRF-3021a. Of course, there still remains the possibility that the anti-tRF is de-repressing the translation of tRF-target mRNAs without affecting the mRNA levels.

### DNA damage following tRF-3021a knockdown

Given the early induction of stress-associated genes (UV_RESPONSE_UP, TNFα signaling via NF-κB) and later enrichment of apoptosis/p53 pathways, we asked whether these transcriptional changes were preceded by activation of a DNA damage response (DDR). We assessed DDR markers at 2 and 4 h following tRF-3021a knockdown. Immunoblotting revealed increased phosphorylation of ATM (p-ATM) and elevated γH2AX levels in tRF-3021a–depleted cells compared with control cells **(Supplementary Fig. 4E-F)**. Taken together, these findings indicate that ATM/γH2AX activation occurs early after tRF-3021a depletion and may represent an upstream response that contributes to the later apoptotic program.

## Discussion

tRNA-derived fragments have been implicated in many diseases [8], but their roles in cancers remain relatively underexplored compared with protein-coding genes, miRNAs, and lncRNAs [6, 37, 38]. We and others have proposed that some tRFs regulate gene expression by entering into Ago containing complexes and repressing target genes by miRNA-like 3′UTR repression mechanism [5, 7, 12, 13, 36, 39]. Consistent with this, tRNA-Leu-CAA–derived tsRNAs (ts-26, tRF-3012a/b) were reported to be reduced in diffuse gliomas and suggested to regulate *RBM43* or *HOXA13*, while tRNA-Cys-GCA–derived tRF-3003a/b were also reduced in gliomas and proposed to repress *VAV2 [40, 41]*. Similarly, two tRNA-Arg–derived tRF-5s were reported to be lower in GBM and to restrain proliferation/migration likely via *S100A11* [38]. Other work has profiled tsRNA landscapes across GBM and LGG [42]. tRFs have also been described in exosomes. For example, exosomal i-tRF-Leu-CAA was proposed as a plasma biomarker and promoted invasion/proliferation (potentially via TPM4/EMT) [43]. More recently, TRMT10A loss was linked to altered tRNA modification and increased tRF-22, promoting vasculogenic mimicry and tumor growth through *MXD1/HIF1A* signaling [44].

In the context of the growing evidence for tRF involvement in cancer biology, our TCGA analysis showed that higher levels of tRF-3009a (tRNA-Leu), tRF-3021a (tRNA-Ala), and tRF-3030a (tRNA-Tyr) associate with poorer survival in LGG. We therefore asked whether these fragments contribute to GBM-relevant phenotypes, but discovered that the phenotypes are similar in multiple cancer lineages. Depletion of each fragment reduced cancer cell invasion and migration, whereas overexpression increased these phenotypes, indicating that these tRF-3s promote cellular behaviors consistent with the infiltrative nature of tumors [45]. Consistent with these invasion/migration phenotypes, we observed reduced SLUG/SNAI2 protein levels upon knockdown of all three tRFs, with the most pronounced effect following tRF-3021a depletion, where SLUG was nearly undetectable. SLUG has been reported to be overexpressed in GBM and to promote mesenchymal/invasive programs [46]. Although multiple transcription factors regulate mesenchymal/invasive programs, the pronounced reduction in SLUG protein upon tRF-3021a knockdown could, in part, contribute to the decreased invasion and migration. The tRF-3 mediated microRNA-like target gene repression cannot explain the repression of the invasive-state genes when the tRF-3 is knocked down. Whether tRF-3021a regulates SLUG directly or indirectly, by affecting SLUG mRNA translation, protein stability or half-life, or upstream signaling, or whether global reductions in protein synthesis following tRF-3021a depletion secondarily lower SLUG levels remains to be determined and should be explored further. Importantly, our early RNA-seq data support suppression of pro-migratory signaling nodes, as pathways typically associated with invasive states, including WNT and TGF-β signaling, were negatively enriched as early as 8 h following tRF-3021a knockdown, both of which can act upstream of mesenchymal transition programs [47].

Among the three candidates, tRF-3021a regulates additional phenotypes beyond invasion/migration and was uniquely required for proliferation, protection from apoptosis, and maintenance of global protein synthesis, suggesting potential relevance as a therapeutic target. Outside glioma, tRF-3021a has been implicated in cancer biology in a context-dependent manner, including reported roles in colorectal cancer immune evasion and cancer progression [48]. In lung cancer, Skeparnias et al. reported that a tRF-3021a species derived from tRNA-Ala-CGC is downregulated and linked to lung cancer progression [49].

Mechanistically, tRF-3021a depletion attenuated global protein synthesis (puromycin or HPG incorporation) across multiple glioma, non-glioma, and normal astrocyte cell types, and translation suppression could be rescued by reintroduction of a tRF-3021a mimic. In contrast, depletion of tRF-3009a, tRF-3030a, and two highly abundant U251 miRNAs (hsa-miR-7-5p and hsa-miR-9-5p) did not reduce global protein synthesis, supporting a specific requirement for tRF-3021a in sustaining translational output.

The sequence specificity of this requirement was further confirmed by different approaches. Rescue of translation required a mimic with unchanged nucleotides in the 5’ fourteen bases corresponding to the tRF. Point mutations in those bases did not abrogate the ability of those mimics from annealing to the LNA, suggesting that the rescue by the tRF mimics was not simply because they annealed to the LNA and titrated it away from other effector nucleic acids or proteins. A single-nucleotide mutant LNA (PM1) retained translation block activity whereas a three-nucleotide mutant LNA (PM3) did not, again showing that even modest sequence perturbations are sufficient to abrogate the function of the LNA. Finally, LNAs targeting either the 5′ end or the anticodon loop of the parental tRNA-Ala did not inhibit translation, suggesting the protein synthesis inhibition was specifically due to the targeting of tRF-3021a and not the parental tRNA. Northern blots and a GCA-codon reporter assay also confirmed that tRNA-Ala levels or decoding activity were unaffected by the anti- tRF-3021a LNA. Together, these data rule out disruption of the parental tRNA-Ala as an explanation for the translational phenotype and establish that the protein synthesis inhibition is specifically due to the targeting and depletion of the tRF-3021a.

Importantly, the earliest phenotypes following tRF-3021a depletion were inhibition of protein synthesis and induction of DNA damage signaling (γH2AX and p-ATM). However, the mechanistic relationship between translational suppression, DNA damage signaling, and the later onset of apoptosis remains unresolved.

Protein synthesis is tightly regulated and essential for cellular homeostasis, and dysregulated translation can reprogram the cellular proteome and contribute to human disease. Translational control can be altered through changes in translation factors (initiation, elongation, and termination), tRNA abundance or function, and regulatory RNAs such as tRFs, collectively shaping production of proteins that support tumor initiation and progression [50, 51]. tRNA-derived fragments (halves and tRFs) can act as both negative and positive regulators of translation [52]. Ivanov et al. showed that stress-activated angiogenin cleaves tRNAs into tiRNAs and that transfection of natural or synthetic tiRNAs is sufficient to inhibit protein synthesis and promote stress granule (SG) assembly even in the absence of eIF2α phosphorylation. Mechanistically, 5′-tiRNAs disrupt the eIF4F complex in a manner requiring a 5′ terminal oligoguanine (TOG) motif [25]. Sobala et al. showed that short 5′-tRFs bearing a conserved 3′-terminal GG motif can repress protein translation in HeLa cells, and that endogenous tRF(Gln) partly co-sediments with polysome fractions, consistent with direct engagement of the translational machinery [53]. In contrast to these translation-repressive tRNA halves and 5′ tRFs, tRF-3021a is a shorter tRF-3 derived from the opposite end of the parental tRNA, and is required to sustain global protein synthesis, suggesting that it may regulate protein synthesis through a distinct mechanism.

Certain tRFs can also enhance translation of specific mRNAs. For example, the LeuCAG3′tsRNA (a tRF-3-like species) binds the CDS and 3′UTR of RPS28 mRNA, remodels its secondary structure, and increases translation of this ribosomal protein transcript to support ribosome biogenesis [26]. In our system, tRF-3021a is required to sustain global protein synthesis, as its knockdown reduces puromycin or HPG incorporation, but the mechanism by which it maintains global translational levels remains unclear.

We examined possible mechanisms by which protein synthesis can be inhibited globally. There is an early decrease in phosphorylation of mTORC1 effectors (p-4E-BP and p-P70S6K) following tRF-3021a depletion and so tRF-3021a may act upstream of mTORC1-dependent translational control. Phosphorylation of 4E-BP reduces its binding to eIF4E, thereby permitting eIF4E–eIF4G assembly and cap-dependent initiation. Accordingly, decreased p-4E-BP could restrain cap-dependent protein synthesis [54], but the extent of decrease in 4E-BP phosphorylation does not seem sufficient to produce the marked global inhibition of protein synthesis that is seen after tRF-3021a knockdown. Paradoxically, tRF-3021a overexpression in A549 lung cancer cells suppresses mTORC1-associated signaling [49]. Another mechanism by which global protein synthesis could be shut down is through the integrated stress response (ISR), in which eIF2α is phosphorylated by one of four upstream kinases (HRI, PKR, PERK, or GCN2) [55], which results in inhibition of protein synthesis by inhibiting eIF2B, impairing its guanine nucleotide exchange factor (GEF) activity, and reducing the eIF2•GTP•Met-tRNAi ternary complex, thereby shutting down global translation initiation [56]. However, in our hands eIF2α phosphorylation was seen much after the inhibition of protein synthesis, and so cannot be the primary cause of protein synthesis inhibition after tRF-3021a knockdown. The ISR activation is a late response, perhaps from stress created by the protein synthesis inhibition, and is known to help either with cellular adaptation or promotion of apoptosis. A hallmark of ISR activation is selective enhancement of ATF4 translation owing to its two uORFs [57], which can induce transcriptional programs including DDIT3 (CHOP) and PPP1R15A (GADD34). Consistent with ISR engagement at 48 h, our RNA-seq shows upregulation of *DDIT3* (CHOP) and *PPP1R15A* (GADD34) together with pro-apoptotic BH3-only transcripts (*BCL2L11*/BIM, *PMAIP1*/NOXA, and *BBC3*/PUMA) and additional stress/DDR-associated genes such as *GADD45G* and *CDKN1A*, aligning with enrichment of p53 and apoptosis pathways in our GSEA analysis. Thus, it is likely that the ISR activation at later time points contributes to downstream stress and apoptotic outcomes. In the astrocytes alone, tRF-3021a knockdown suppressed global protein synthesis, but enhanced the incorporation of puromycin in short polypeptide chains. This suggests that at least in astrocytes the global protein synthesis inhibition may stem from a block to polypeptide chain elongation rather than inhibition of protein synthesis initiation. The mechanism for such a block to elongation is unclear, but it is unlikely to be due to depletion of the parental tRNA of tRF-3021a, because (a) we do not see a decrease in the level of that tRNA, (b) antisense to other tRFs (and their parental tRNAs) do not decrease protein synthesis and (c) we have experimentally shown that levels and activity of the parental tRNA-Ala are not decreased on an alanine-codon reporter by the anti-tRF-3021a LNA.

To better understand early events, inspection of the 8 h Reactome GSEA revealed enrichment of pathways related to translation initiation and elongation, as well as response to amino acid/tRNA deficiency. The amino acid deficiency response is canonically triggered when uncharged (deacylated) tRNAs accumulate and activate GCN2 at the ribosome [58]. Although we did not detect early eIF2α phosphorylation, a downstream readout of canonical ISR activation, enrichment of this pathway suggests an early transcriptional signature of amino acid limitation/translation stress. Interestingly, despite the translation repression, we see an enrichment of translation associated mRNA sets (**Supplementary Fig. 4B**) which might reflect a compensatory transcriptional feedback response to translation inhibition. Supporting this idea, in yeast, compensatory induction of ribosome biogenesis related mRNAs has been reported in response to global translation inhibition [59]. Together, these findings support early disruption of translational homeostasis following tRF-3021a depletion, although the mechanism by which tRF-3021a suppresses global translation remains to be determined.

In parallel with these translation-linked signatures, Hallmark GSEA at 8 h revealed positive enrichment of oxidative phosphorylation (OXPHOS) gene sets. Whether this transcriptional signature reflects increased oxygen consumption or altered mitochondrial activity remains to be determined. Nevertheless, consistent with the possibility that tRFs can influence mitochondrial programs, Geng et al. reported that tRF-3009a is required for IFN-α–induced oxidative phosphorylation, ATP production, and ROS generation in lupus CD4⁺ T cells [60]. TNFα signaling via NF-κB enriched at 8 h and remained elevated at 48 h. Leading-edge genes at 8 h included stress and checkpoint regulators (GADD45G, GADD45B, and CDKN1A). The GADD45 family encodes small, stress-inducible proteins that lack intrinsic enzymatic activity and therefore function primarily through protein–protein interactions with regulators of checkpoint control and DNA repair such as p21, PCNA, and CDK1/cyclinB1. Consistent with their established roles in cellular injury responses, GADD45 proteins are rapidly induced following DNA damage and can promote cell-cycle arrest, DNA repair, and apoptosis [61, 62]. Importantly, GADD45 proteins can couple cellular stress to apoptosis by binding and activating the stress-responsive MAPKKK MTK1/MEKK4, triggering downstream p38 and JNK signaling and promoting apoptotic outcomes [39].

Because DNA damage response (DDR) signaling was detectable as early as 4 h (γH2AX and p-ATM), these early transcriptional changes may reflect activation of a stress program downstream of early DDR engagement. Persistent activation of these pathways may contribute to the later enrichment of apoptotic signatures and commitment to cell death. Supporting the concept that tRFs can indirectly modulate H2AX phosphorylation, Maute et al. showed that CU1276 (a tRF-3b) represses *RPA1* at the translational level and sensitizes lymphoma cells to etoposide-induced DNA damage, as measured by increased γH2AX [12]. How tRF-3021a protects cells from DNA damage and apoptosis remains unclear, and whether DDR activation and translational inhibition occur in parallel or are mechanistically linked will require further investigation.

We and others have reported that tRF-3s can be incorporated into Argonaute-containing silencing complexes and repress protein and mRNA levels of genes harboring complementary target sites, following microRNA-like seed sequence rules [4, 12, 13, 63, 64]. Surprisingly, depletion of tRF-3021a did not result in robust de-repression of predicted targets, nor did any candidate targets explain the magnitude of the phenotypic changes observed following depletion (data not shown). These findings suggest that the mechanism by which tRF-3021a promotes global protein synthesis and limits apoptosis is unlikely to be driven primarily by canonical microRNA-like gene silencing.

In summary, our study identifies three tRF-3s whose higher expression is associated with poorer outcome in LGG in a univariate prioritization analysis and demonstrates that these fragments promote cancer cell invasion and migration. We further show that tRF-3021a is uniquely required to sustain proliferation, limit apoptosis, and maintain normal global protein synthesis, in many different cancer cell lineages. The translational inhibition occurs early and precedes apoptosis. Some DNA damage and signs of oxidative stress also appear early. Multiple experiments, including rescue by mutant mimic, translational block by mutant LNAs, targeting the parental tRNA by other LNAs, and a GCA-codon reporter assay, demonstrate that the translational phenotype is specifically due to the loss of the tRF-3021a fragment and not to disruption of the parental tRNA-Ala. Together, these findings establish tRF-3021a as a regulator of cancer cell biology, specifically translational homeostasis and motivate future studies to define the direct molecular interactors and mechanistic steps linking tRF-3021a to translational control. Elucidating this pathway may reveal opportunities to exploit tRF-3021a for potential therapeutic vulnerability.

## Acknowledgement

This work was supported by R01 GM146756 (to AD) and R00 CA259526 (to ZS). We thank all members of the Dutta and Su labs for help and advice

ChatGPT (OpenAI) was used to assist with editing and polishing manuscript, and all content was reviewed and validated by the authors.

## Supplementary data

**Supplementary Figure 1.**
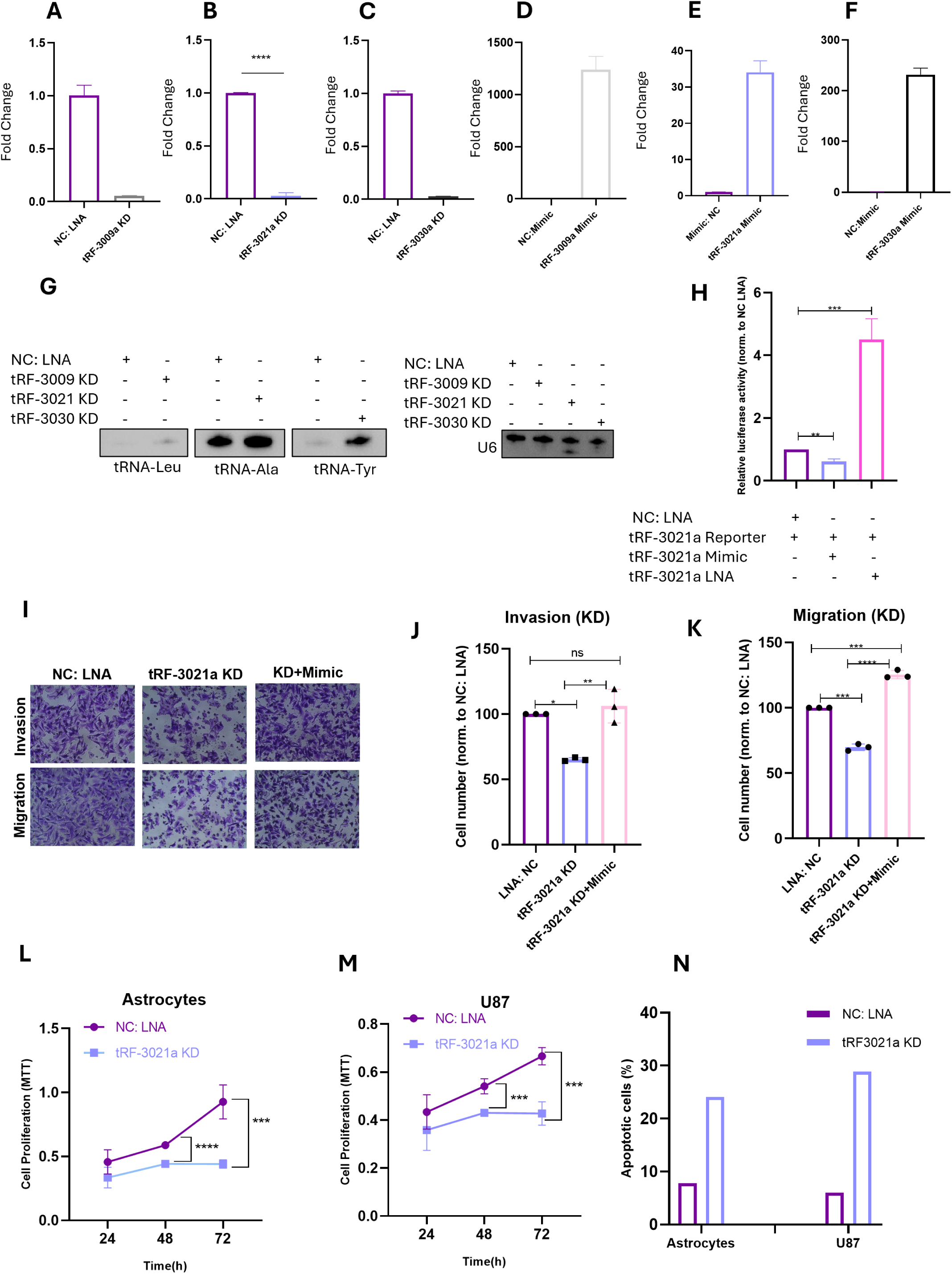
Validation of tRF-3 knockdown and overexpression and additional functional assays supporting tRF-3021a activity. (A–C) qRT-PCR validation of LNA-mediated knockdown (KD) of tRF-3009a (A), tRF-3021a (B), and tRF-3030a (C) in U251 cells compared with negative-control (NC) LNA. Expression was normalized to NC LNA. (D–F) qRT-PCR validation of mimic-mediated overexpression (OE) of tRF-3009a (D), tRF-3021a (E), and tRF-3030a (F) in U251 cells compared with NC mimic. Expression was normalized to NC mimic. (G) Northern blot analysis of parental tRNAs in U251 after KD of tRF-3009a, tRF-3021a, or tRF-3030a compared with NC LNA. U6 was used as a loading control. (H) Dual luciferase reporter assay in U251 cells using a tRF-3021a reporter. Reporter activity was normalized to the NC LNA reporter condition. (I) Representative Transwell images showing invasion and migration of U251 cells after 12 h after tRF-3021a KD and rescue with tRF-3021a mimic. (J–K) Quantification of invasion (J) and migration (K) in the rescue experiment. Cell numbers were normalized to NC LNA, which was set to 100%. (L–M) MTT assays in normal astrocytes (L) and U87 glioma cells (M) after tRF-3021a KD compared with NC LNA at the indicated time points. (N) Annexin V-positive cells in normal astrocytes and U87 glioma cells after tRF-3021a KD compared with NC LNA. Data are presented as mean ± SD. For qRT-PCR validation, tRF-3021a KD and OE were analyzed from independent biological replicates, whereas tRF-3009a and tRF-3030a KD and OE validation panels show technical qPCR replicates. Statistical significance was determined by unpaired two-tailed t-test for qRT-PCR and luciferase reporter comparisons, one-way ANOVA with multiple-comparisons testing for the Transwell rescue experiments, and two-way ANOVA with multiple-comparisons testing for MTT time-course experiments. Panel N was performed once. ns, not significant, *P < 0.05, **P < 0.01, ***P < 0.001, ****P < 0.0001.

**Supplementary Figure 2.**
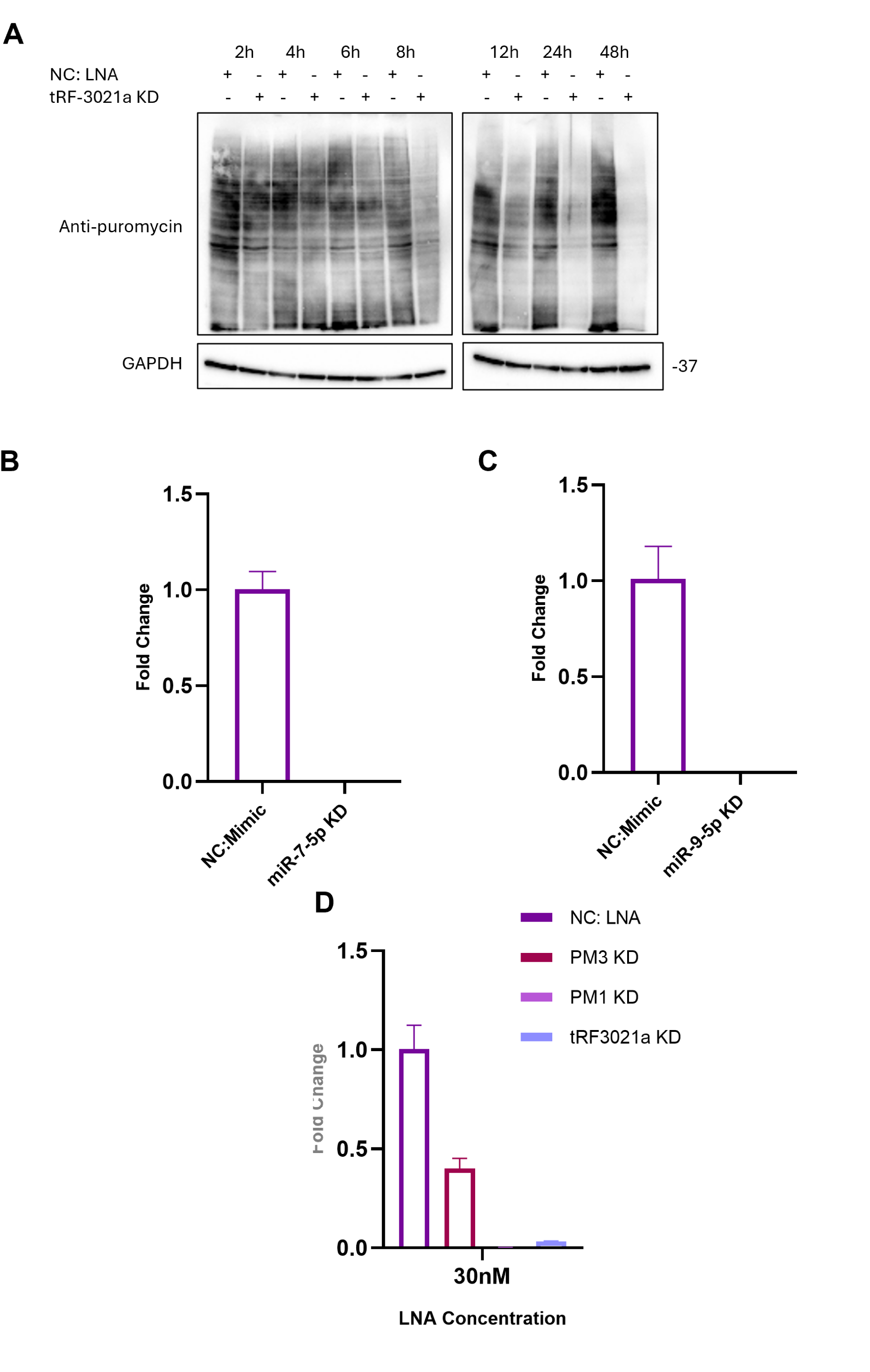
Representative puromycin blot for time-course quantification, miRNA knockdown validation, and tRF-3021a levels after tRF-3021a KD and point-mutated LNAs. (A) Representative puromycin incorporation immunoblot corresponding to the quantification shown in Fig. 5C. GAPDH was used as a loading control. (B–C) qRT-PCR validation of miR-7-5p knockdown (B) and miR-9-5p knockdown (C) compared with negative-control (NC) LNA. Expression was normalized to NC LNA. (D) qRT-PCR analysis of tRF-3021a levels after transfection with tRF-3021a LNA or point-mutated tRF-3021a LNAs. PM1 decreased tRF-3021a levels to a similar extent as the tRF-3021a LNA, whereas PM3 showed a weaker reduction in tRF-3021a levels. Data in panels B–D are shown as mean ± SD from three technical replicates.

**Supplementary Figure 3.**
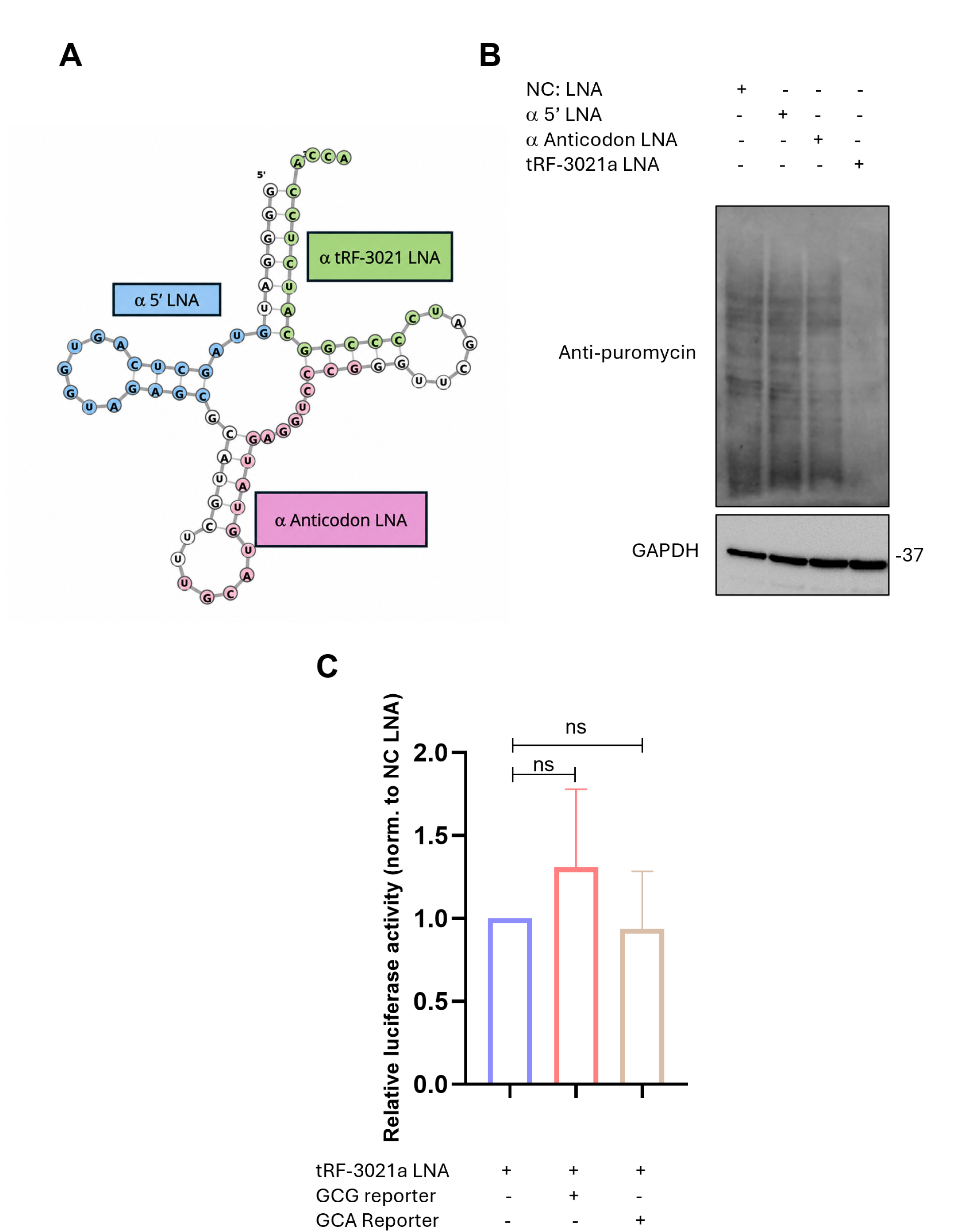
tRF-3021a-mediated translation inhibition is not caused by disruption of parental tRNA-Ala. (A) Schematic showing the positions of LNAs targeting different regions of parental tRNA-Ala-TGC, including a 5′ LNA, an anticodon LNA, and the tRF-3021a LNA. (B) Puromycin incorporation assay in U251 cells transfected with negative-control (NC) LNA, 5′ LNA, anticodon LNA, or tRF-3021a LNA. GAPDH was used as a loading control. (C) GCG- and GCA-codon reporter assay in U251 cells after tRF-3021a KD. Reporter activity was normalized to the tRF-3021a KD + empty-vector condition. Data are presented as mean ± SD from independent biological replicates unless otherwise indicated. Statistical significance was determined by unpaired two-tailed t-test for the codon-reporter assay. Immunoblots are representative experiments unless otherwise indicated. ns, not significant, *P < 0.05, **P < 0.01, ***P < 0.001, ****P < 0.0001.

**Supplementary Figure 4.**
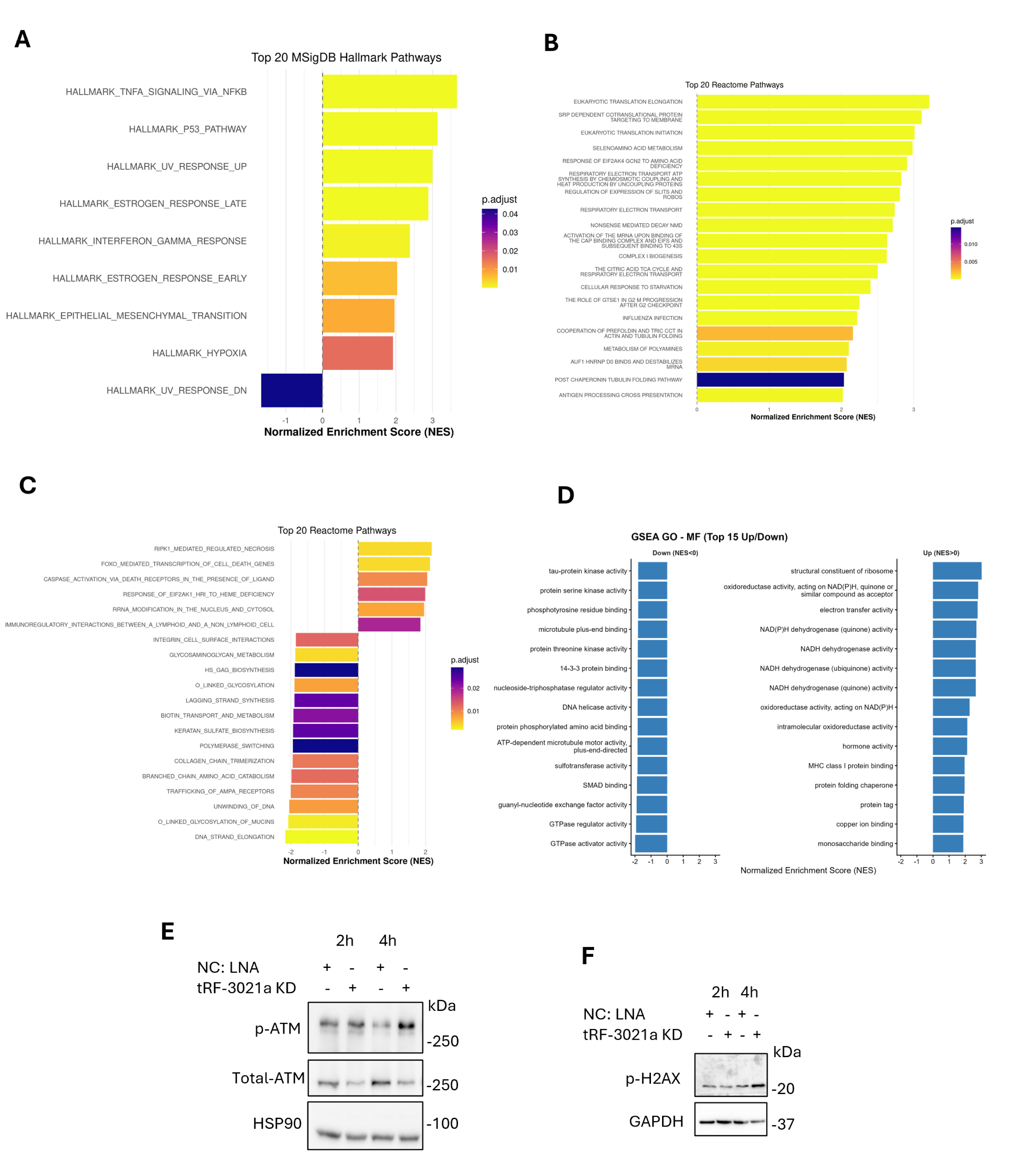
Shared Hallmark Enrichment, Reactome Changes, and Early DNA Damage Activation following tRF-3021a Knockdown. (A) MSigDB Hallmark pathway enrichment analysis of genes commonly regulated in U251 cells after LNA-mediated knockdown (KD) of tRF-3021a compared with negative-control (NC) LNA at 8 h and 48 h. (B–C) Reactome pathway enrichment analysis of genes altered after tRF-3021a KD compared with NC LNA at 8 h (B) and 48 h (C). (D) Gene Ontology molecular function enrichment analysis showing the top downregulated and upregulated pathways after tRF-3021a KD. Normalized enrichment scores are shown, and adjusted P values are indicated by color. (E) Immunoblot analysis of phosphorylated ATM and total ATM in U251 cells after tRF-3021a KD compared with NC LNA at 2 h and 4 h. HSP90 was used as a loading control. (F) Immunoblot analysis of phosphorylated H2AX in U251 cells after tRF-3021a KD compared with NC LNA at 2 h and 4 h. GAPDH was used as a loading control. GSEA results are presented as normalized enrichment scores with adjusted P values. Immunoblots are representative experiments unless otherwise indicated.

**Supplementary Table 1. Differential gene expression analysis following tRF-3021a knockdown**.

This table provides the complete edgeR statistical results for gene expression comparisons between tRF-3021a knockdown (KD) and negative-control (NC) LNA in U251 cells.

8 h sheet: edgeR results for the 8 h comparison.

48 h sheet: edgeR results for the 48 h comparison.

8 h FDR sheet: genes with FDR < 0.05 at 8 h.

8 h up sheet: upregulated genes with FDR < 0.05 and |log2 fold change| ≥ 1 at 8 h.

8 h down sheet: downregulated genes with FDR < 0.05 and |log2 fold change| ≥ 1 at 8 h.

48 h FDR sheet: genes with FDR < 0.05 at 48 h.

48 h up sheet: upregulated genes with FDR < 0.05 and |log2 fold change| ≥ 1 at 48 h.

48 h down sheet: downregulated genes with FDR < 0.05 and |log2 fold change| ≥ 1 at 48 h.

